# Asymmetric Division in a Two-Cell-Like State Rejuvenates Embryonic Stem Cells

**DOI:** 10.64898/2026.01.07.697614

**Authors:** Xinyi Wang, Hong Fu, Qingyang Sun, Boyan Huang, Zhe Xu, Xuzhao Zhai, Chuncao Deng, Laru Peng, Mengdan Zhang, Tianran Peng, An Gong, Jiasui Liu, Zhengzhi Zou, Jiekai Chen, Guangming Wu, Man Zhang, Mingwei Min

## Abstract

A fundamental question in biology is whether cellular aging is inevitable. Embryonic stem cells (ESCs) challenge this paradigm as rare normal cells capable of indefinite in vitro passage. However, the mechanisms underlying ESC lineage immortality remain unresolved. Using long-term live-cell imaging to follow fates of single ESCs, we show here that ESC lineage renewal is achieved through sporadic entry into a state characterized by the expression of two-cell embryo-specific markers. During this state, cells undergo asymmetric divisions, segregating accumulated DNA damage into one daughter lineage destined for elimination, while producing a second lineage that reverts to the pluripotent state. Importantly, the latter lineage exhibits signs of rejuvenation, including reduced DNA damage and enhanced chimeric efficiency. These findings underscore the crucial role of asymmetric cell division in maintaining the long-term health of the ESC lineage against mounting damage within individual cells, a potential model for studying cellular aging and rejuvenation in mammalian cells.

## Introduction

The lineage of all life today traces back to an ancestor 3.8 billion years ago. While the overall life lineage appears to be immortal, individual organisms succumb to aging and mortality. This duality is often attributed to the division of labor between germline and soma. In this framework, the germline perpetuates the lineage, whereas the soma accumulates damage and is ultimately disposable^1,2^. Most somatic cells exhibit progressive functional decline during proliferation – a phenomenon classically attributed to telomere attrition, DNA damage, and epigenetic drift. Embryonic stem cells (ESCs) however, defy this trajectory, achieving lineage immortality through indefinite in vitro proliferation while maintaining its function, a long-sought property in regenerative medicine. Specifically, these long-term passaged ESCs maintain a normal karyotype and the capacity to contribute to chimeric animals^3,4^. While telomere maintenance mechanisms in ES cells are well characterized^5–7^, it remains unclear how these cells evade the cumulative burden of molecular damage that limits the lifespan of other cell types.

A clue lies in the transient activation of programs associated with the two-cell (2C) embryo stage. ESCs occasionally enter a 2C-like state marked by the expression of genes such as *MERVL* and *Zscan4*^5,8^. Intriguingly, knockdown of *Zscan4* or depletion of MERVL expressing cells induces culture crisis of ESCs^5,9^, while *Zscan4* overexpression enhances pluripotency^10^, suggesting the spontaneous 2C-like state may serve as a ‘fountain of youth’ for the ESC lineage. Paradoxically, 2C-like cells exhibit increased DNA damage^11^, underscoring a gap in our understanding of how the transient 2C-like states might contribute to long-term lineage renewal of ESCs.

We propose two non-exclusive hypotheses: 2C-like cells either employ unusually efficient biochemical repair mechanisms to counteract cellular aging, or sort away damaged components. Using long-term live-cell imaging, we tracked the fate and the damage status of individual ESCs spontaneously entering the 2C-like state. Our data support the latter hypothesis. Upon entering the 2C-like state, cells undergo asymmetric divisions, producing two distinct progenies: one lineage enriched for DNA damage and prone to cell death, while the other that reacquires pluripotency with reduced damage. Functional assays confirm that the latter lineage exhibits enhanced chimeric contribution in developing embryos, whereas blocking entry into the 2C-like state compromises ESC self-renewal. These findings position asymmetric division within the 2C-like state as a critical quality-control mechanism, enabling ESCs to sustain lineage immortality despite spontaneous DNA damage. Our work provides a framework for understanding how stem cells safeguard genetic perpetuity of the lineage – a paradigm with implications for regenerative medicine, aging, and cancer biology.

## Results

### 2C-like cells undergo asymmetric divisions

To trace the fate of 2C-like cells, we established a mouse embryonic stem cell (mESC) line expressing a MERVL promoter-driven GFP reporter (MERVL::GFP) to mark the 2C-like state^8^, and H2B-mIFP to track cells. The MERVL::GFP signal strongly correlates with endogenous MERVL-Gag protein staining (Figure S1A). Using time-lapse imaging, we followed the lineages of cells entering the 2C-like state (Figure 1A). Surprisingly, cells displaying a strong MERVL::GFP signal predominantly underwent cell death (Figures 1B-1C, S1B and Video S1), contrary to the hypothesis that the 2C-like state promotes the renewal of mESCs. Quantification of lineage survival indicated that approximately 40% of the MERVL::GFP^+^ cells died within 72 hours (Figure 1D). The remaining living MERVL::GFP^+^ cells exhibited diminished GFP intensity (Figures 1E-1F), and progressively transitioned into a MERVL::GFP^−^ state within a few generations (Video S2). We denote these two fates as 2C-death and 2C-survived, respectively.

**Figure 1.**
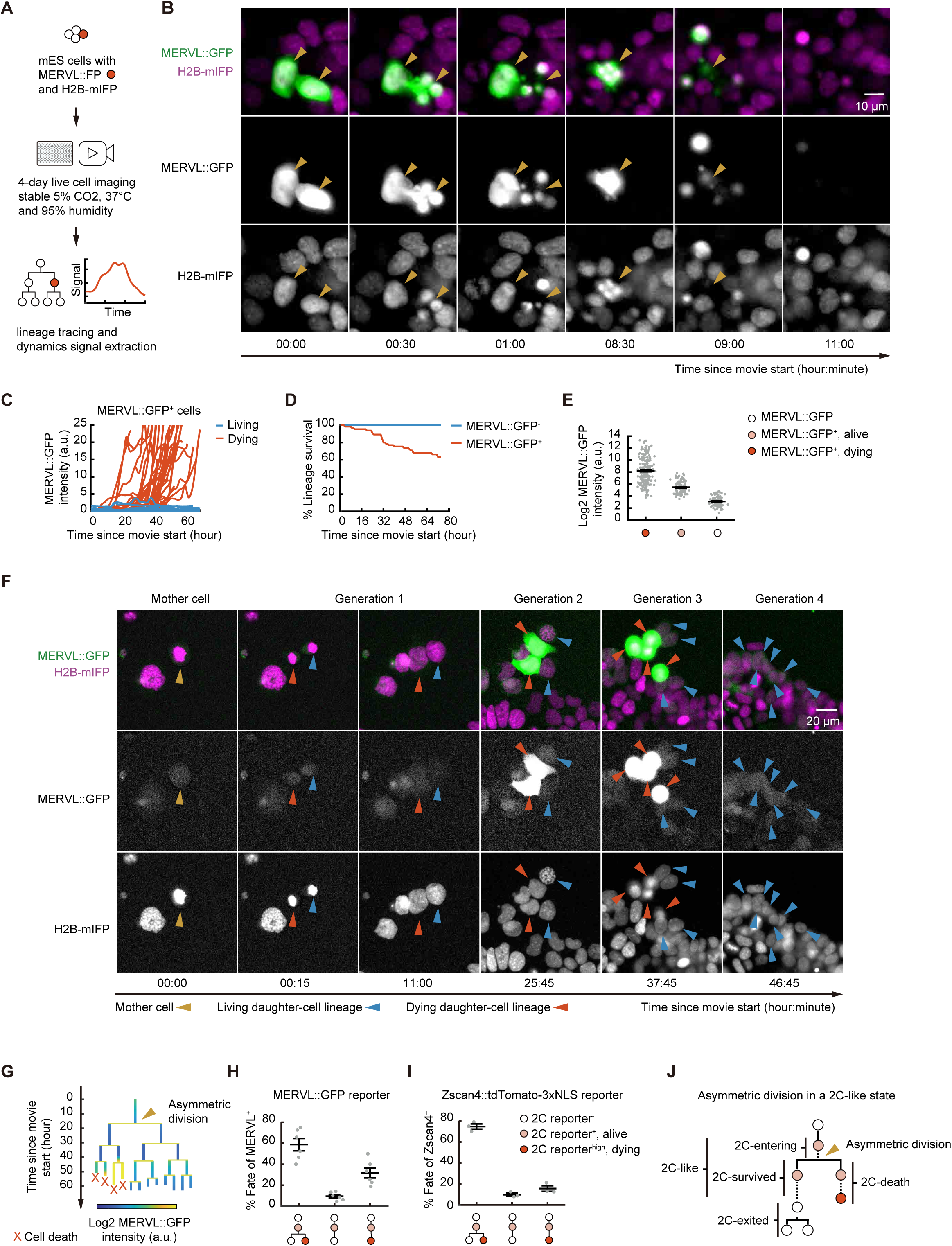
2C reporter^+^ cells generate asymmetric daughter-cell fates. (A) Experimental scheme of live cell imaging experiments. mES cells were cultured in 96-well plate with stable 5% CO_2_, 37°C and 95% humidity. The fluorescent signal of H2B-mIFP was used to track cell lineage and MERVL::FP to monitor MERVL status. (B) Examples of MERVL::GFP^+^ cell death, indicated by the condensation or fragmentation of H2B-mIFP signal without cell division, and the subsequent disappearance of MERVL::GFP signal within 1-2 frames (15-30 minutes). Scale bar, 10 μm. See also Figure S1B for other examples and Video S1 for the full process. (C) Dynamics of GFP intensity in MERVL::GFP^+^ cells, grouped by their living or dying fates determined by the end of the movie (n = 50 in each group). (D) Lineage survival of MERVL::GFP^+^ (n = 65) and MERVL::GFP^−^ (n = 85) cells, plotted as a Kaplan-Meyer curve. For MERVL::GFP^+^ cells, lineage was tracked from the moment of MERVL::GFP^−^ to MERVL::GFP^+^ transition and classified as “death” if all progenies died by the end of the movie, or “alive” if any progeny survived. For MERVL::GFP^−^ cells, lineages that remained MERVL::GFP^−^ throughout the entire movie were tracked and assessed identically. (E) MERVL::GFP intensity of MERVL::GFP^−^ (n = 95), living (n = 103) and dying (n = 190) MERVL::GFP^+^ cells. Each data point represents the average MERVL::GFP intensity over one cell cycle. (F) An example of a MERVL::GFP^+^ cell producing daughter cells with distinct fates. The images, from left to right in the temporal order from a timelapse movie, show the fates of all progenies from a MERVL::GFP^+^ cell. The mother cell (yellow arrowhead), exhibiting low but above-background MERVL::GFP expression, divided into two daughter cells (column 1-2). One daughter generated a lineage with increasing levels of MERVL::GFP (column 3-4) and all cells died by 46 hours (tracked by red arrowheads). The other daughter produced a viable lineage with diminishing MERVL::GFP levels (tracked by blue arrowheads). Scale bar, 20 μm. See Video S2 for another example. (G) Lineage tree of a MERVL::GFP^+^ cell. The cell entered a low-expression MERVL::GFP^+^ state (0-18 hour). One of its daughters showed strongly increasing MERVL::GFP levels (starting around 42 hour), with all progenies died by 60 hour (marked by crosses). The other daughter produced a lineage with all progenies alive. The arrowhead indicates an asymmetric division, assigned retrospectively. The heat colors indicate the fluorescence intensity of MERVL::GFP. (H, I) Percentage of the different daughter-cell fates of ESCs spontaneously transitioning into the 2C-like state, marked by either MERVL::GFP^+^ (H) or Zscan4::tdTomato^+^ cells (I); each point represents an experiment. (J) A schematic to depict subpopulations of 2C-like cells. The subpopulations are defined in live cell movies with the lineage dynamics. The arrowhead indicates the asymmetric division of daughter-cells fates. Error bars indicate mean ± SEM (E, H-I). See also Figure S1 and Videos S1-S3.

Are the two fates independent or interconnected? Upon entering the MERVL::GFP^+^ state, about 30% cells generated only 2C-death fate, while 10% generated only 2C-survived fate. The 2C-survived-only cells generally expressed very low levels of 2C reporter, suggesting that their 2C-like program is not fully active. About 60% cells gave rise to two daughter cells with distinct lineage fates – one 2C-death and the other 2C-survived (Figures 1F-1H, S1C, and Video S2). The prevalence of coupled cell fates far exceeds that would be expected by random chance: Given that approximately 1% of mESCs are MERVL^+^ at any given time^8^, the probability of a cell generating 2C-survived or 2C-death fate can be derived as P(2C-survived) = *1%* × (*0.1* + *0.6*) = *7* × *10*^−*3*^ and P(2C-death) = *1%* × (*0.3* + *0.6*) = *9* × *10*^−*3*^. If the two fates are independent, the chance of them occurring in the same lineage would be P(2C-survived) × P(2C-death) = 63× *10*^−*6*^, two orders of magnitude less than the observed probability P(2C-survived, 2C-death) = *1%* × *0.6* = *6* × *10*^−*3*^. Hence, the 2C-death and 2C-survived lineages are likely generated by the same progenitor cell, suggesting that the cells entering the 2C-like state can undergo asymmetric division. This observation is further corroborated by cells expressing other 2C reporters, MERVL::tdTomato or Zscan4::tdTomato (Figures 1I, S1A, S1D-S1E, and Video S3), and by cells cultured in the 2i/LIF condition that maintains a naïve ground state^12^ (Figure S1E).

The phenotype of asymmetric progeny fates is reminiscent of budding yeast divisions, where the aged mother cell succumbs to death after a limited number of divisions, and the young daughter cell starts anew with an age reset to zero^13^. Asymmetric division in yeast segregates damaged cellular components from newly synthesized ones^14,15^, providing a mechanism for lineage renewal at the cost of individual aging. We hypothesize that mESCs may adopt a similar strategy for lineage renewal through asymmetric divisions during the spontaneous 2C-like state. Our hypothesis predicts the following: 1) cells entering the 2C-like state (thereafter referred to as ‘2C-entering’ cells, Figure 1J) exhibit signs of aging, marked by elevated cellular damage; 2) cells in the 2C-like state asymmetrically allocate damaged components between the two progenies, which dictates their distinct fates; 3) the 2C-survived lineage that prevails through the asymmetric division experiences functional rejuvenation. We tested all the three predictions in this study.

### 2C-like cells show signs of aging

A prominent feature of aging cells is lengthening of the cell cycle^16^. Indeed, MERVL::GFP^+^ cells took longer to progress through the cell cycle (Figure 2A), consistent with previous reports with fixed cell data^17–20^. Cell cycle lengthening is particularly pronounced in the 2C-entering and 2C-death subpopulations (Figure 2B). Notably, a portion of 2C-death cells undergo prolonged cell cycle arrest before death (Figure S2A). Using a cell-cycle phase reporter, PIP-FUCCI^21^ (Figure S2B), we found that the cell cycle lengthening can be attributed to the lengthening of S and G2/M phase (Figure 2C), consistent with the elevated replication stress reported before^22^.

**Figure 2.**
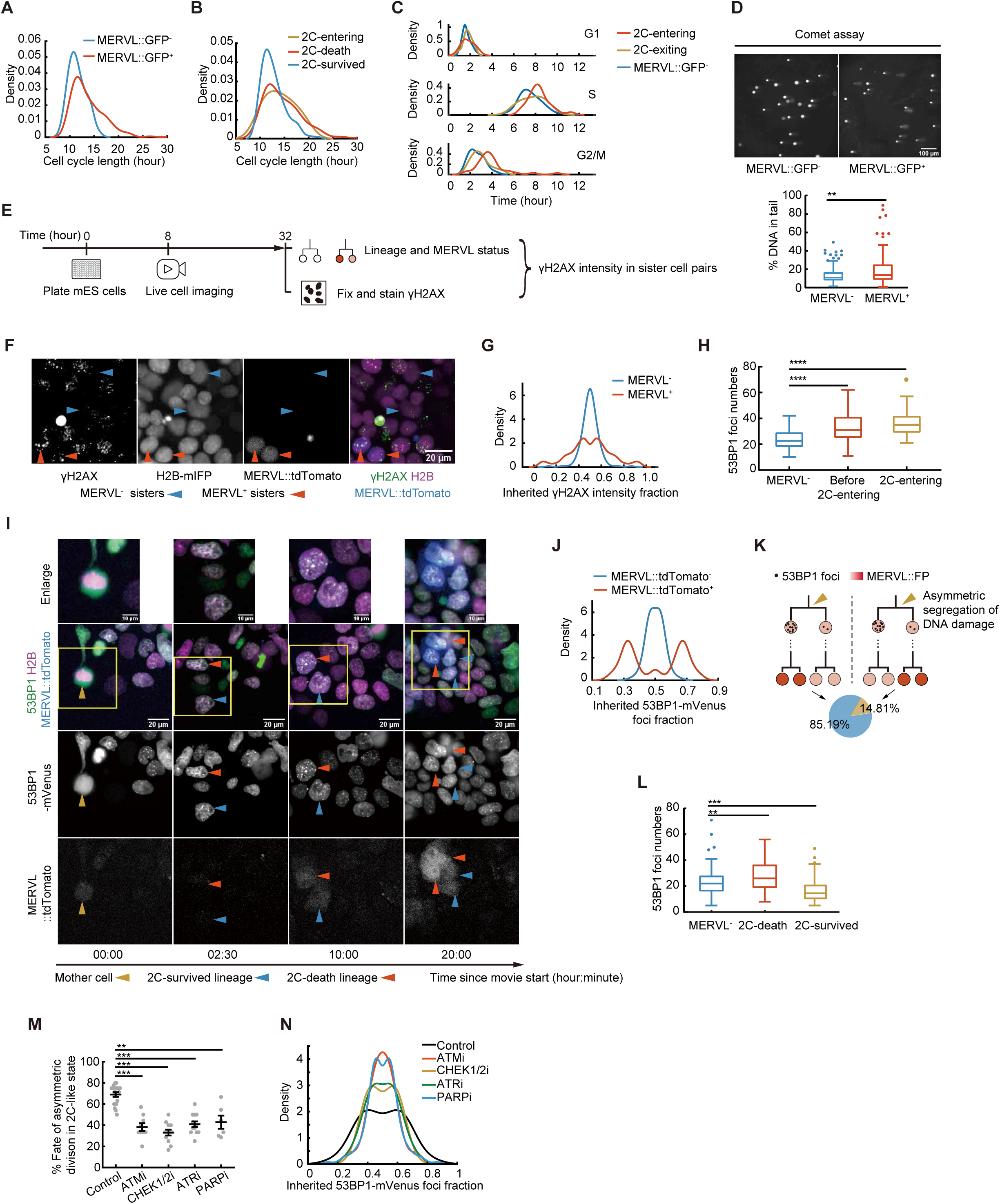
Asymmetric division of DNA damage in 2C reporter^+^ cells. (A) Cell cycle length of MERVL::GFP positive (n = 269) and negative (n = 100) cells, measured as the time between two consecutive mitoses marked by the segregation of H2B-mIFP in live cell imaging experiments. *p* < 0.0001. (B) Cell cycle length of 2C-survived (n = 150), 2C-entering (n = 53, *p* < 0.01), 2C-death (n = 125, *p* < 0.0001) cells among the MERVL::GFP^+^ lineages that undergo asymmetric divisions. (C) The length of cell cycle phases in the 2C-entering (G1: n = 33; G2: n = 37; S: n = 23), 2C-exiting (G1: n = 33; G2: n = 34; S: n = 26) and MERVL::GFP^−^ (G1: n = 106; G2: n = 92; S: n = 80) cells, determined by PIP-FUCCI-mCherry and H2B-mIFP reporters. See more details in Figure S2B. The 2C-entering cells exhibit significant lengthening of the S phase (*p* < 0.01) and G2/M phase (*p* < 0.0001) compared to MERVL::GFP^−^ cells. (D) Comet assay of sorted MERVL::GFP^+^ and MERVL::GFP^−^ cells. Top, representative images; bottom, quantification of the tail DNA fraction (n = 128 for each group). Scale bar, 100 μm. (E) Experiment design of (F) and (G). Live-cell imaging was performed to obtain cell lineage and MERVL status, followed by fixation and anti-γH2AX staining at the endpoint for comparing γH2AX intensity between sister cells. (F) Representative γH2AX staining in MERVL::tdTomato^+^ (red arrowheads) and MERVL::tdTomato^−^ (blue arrowheads) sisters. Scale bar, 20 μm. (G) Density distribution of the inherited γH2AX fraction between 2C-entering MERVL::tdTomato^+^ or MERVL::tdTomato^−^ sisters: γH2AX intensity of each sister divided by the sum of the pair (n = 97 pairs for each group). *p* < 0.0001. (H) 53BP1-mVenus foci numbers in 2C-entering (n = 56), before 2C-entering (one cell cycle before the 2C reporter lighting up, n = 53) and MERVL^−^ cells (n = 56). (I) A representative division of 53BP1-mVenus foci in MERVL^+^ cell lineage. The daughter cell inherited more 53BP1 foci (column 1-3) became 2C-death lineage with higher MERVL::tdTomato intensity (column 3-4). Yellow arrowhead, MERVL^+^ mother cell; red arrowheads, 2C-death lineage; blue arrowheads, 2C-survived lineage. Scale bars, 20 μm. Top row, enlarged images with scale bars 10 μm. See Video S4 for the full progress. (J) Density distribution of the inherited 53BP1-mVenus foci fraction: foci number of each sister divided by the sum of the pair (n = 54 pairs for each condition). *p* < 0.0001. (K) A schematic to depict the two correlation modes between asymmetric segregation of 53BP1 foci number and asymmetric division of MERVL^+^ cells. Left, Cells inheriting more 53BP1 foci numbers adopt the 2C-death fate; right, cells inheriting more 53BP1 foci numbers adopt the 2C-survived fate. The proportion of the two is shown in the pie chart. (n = 54, *p* = 0.008) (L) 53BP1 foci numbers in MERVL^−^ (n = 108), 2C-death (n = 54) and 2C-survived daughter cells (n = 54). (M) Percentage of asymmetric division in 2C-like cells when DNA damage response is inhibited by ATM, CHEK, ATR or PARP inhibitors (ATMi, n = 9; CHEKi, n = 12; ATRi, n = 13; PARPi, n = 6; control, n = 18). (N) The density distribution of inherited 53BP1-mVenus foci fraction in daughter cells under the inhibition of DNA damage response: foci number of each sister divided by the sum of the pair. Control (n = 118 pairs), ATMi (n = 31 pairs, *p* < 0.01), CHEK1/2i (n = 31 pairs, *p* < 0.05), ATRi (n = 31 pairs, *p* < 0.05), PARPi (n = 84 pairs, *p* < 0.001). Error bars indicate mean ± SEM (M). Mann–Whitney U test (A-D, H, L-M); Kolmogorov-Smirnov test (G, J, N); permutation test (K). See also Figures S2-S3 and Video S4.

This result implies that 2C-like cells may contain elevated DNA damage, another aging-related feature^23^. Indeed, MERVL::GFP^+^ cells show increased levels of phospho-p53, a marker of active DNA damage response (Figure S2C). An unbiased transcriptomic examination by single cell RNA sequencing confirms activation of the p53 pathway: direct p53 targets are enriched in genes upregulated in MERVL::GFP^+^ cells compared to MERVL::GFP^−^ cells (*p* = 1.77×10^−6^, hypergeometry test, Figures S2D-S2E). Moreover, the enrichment is even more significant in the cluster 3 with the highest 2C markers expression (*p* = 2.46×10^−13^, hypergeometry test, Figures S2F-S2H), likely corresponding to the 2C-death subpopulation. Using comet assays, we found that sorted MERVL::GFP^+^ cells show a higher fraction of DNA in the comet tails, suggesting that they contain more physical DNA breaks (Figure 2D). Inducing DNA damage by Aphidicolin (Aph), Mitomycin C (MMC) or Hydroxyurea (HU) increases the fraction of MERVL^+^ cells in a dose-dependent manner (Figure S2I), consistent with previous reports^11^. Taken together, our results indicate that 2C-like cells show more signs of DNA damage, which can be a key driver of entry into the 2C-like state.

We further tested if other types of cellular damage promote entry into 2C-like state. Inducing oxidative stress by 2-Methoxyestradiol (2-MeOE2) or osmotic stress by NaCl, sucrose or polyethylene glycol (PEG) increases the fraction of MERVL^+^ cells (Figure S2J), consistent with previous report^24^, whereas endoplasmic reticulum (ER) stress does not significantly alter the percentage of MERVL^+^ cells (Figures S2K-S2L). These results indicate that the entry of the 2C-like state can be driven by certain types of cellular damage.

### DNA damage is asymmetrically segregated in the 2C-like state

Can cellular damage be asymmetrically separated in the 2C-like state? The intensity of γH2AX, a marker of DNA damage, tends to show a broader distribution in MERVL^+^ cells compared to that in MERVL^−^ cells (Figure S3A). When DNA damage is exogenously induced, this trend becomes more pronounced in MERVL^+^ cells, and the distribution appears bimodal (Figure S3A). These observations align with the hypothesis that damaged DNA may be divided asymmetrically in MERVL^+^ cells. To directly test this hypothesis, we sought to investigate the partitioning of damaged DNA between sister cells. We used time-lapse imaging of mESCs with a nuclear marker to trace their divisions and a MERVL reporter to track their MERVL status. At the end of the movie, we fixed the cells and stained them with the γH2AX antibody to assess DNA damage in MERVL^+/−^sister cell pairs (Figure 2E). Notably, asymmetric division of γH2AX is much more prevalent between MERVL^+^ sisters divided from 2C-entering cells compared to MERVL^−^ones (Figures 2F-2G), indicating that damaged DNA tends to be asymmetrically segregated in the 2C-like state.

To test whether asymmetric segregation of damaged DNA correlates with 2C-like cells fate, we introduced a live cell reporter of DNA damage – the tandem Tudor domain of 53BP1 fused to the fluorescent protein mVenus^25^ into an mESC line with MERVL reporter to monitor their temporal dynamics. 53BP1-mVenus forms foci that co-localize with γH2AX foci (Figure S3B), and the foci number increases after DNA damage induction (Figure S3C), indicating that 53BP1-mVenus foci faithfully represent DNA damage. Like γH2AX, the distribution of 53BP1-mVenus foci number appears broader in MERVL^+^ cells than that in MERVL^−^ cells (Figure S3D). In the live cell imaging, both 2C-entering cells and cells one cell cycle before lighting up their 2C reporter contain more 53BP1-mVenus foci than the rest of MERVL^−^ cells (Figure 2H), suggesting that DNA damage precedes entry into the 2C-like state. We further traced the segregation of DNA damage. Importantly, 53BP1-mVenus foci can be asymmetrically divided in 2C-entering cells with weak MERVL::tdTomato signals (Figure 2I, column 1 to 3), more often than in MERVL^−^cells (Figure 2J). Asymmetric segregation of DNA damage occurs well before differences in 2C reporter intensity become apparent between daughter cell lineages (Figure 2I and S3E), suggesting that DNA damage asymmetry is likely a cause rather than a consequence of their distinct cell fates. The progenies inheriting fewer 53BP1 foci were bias towards the 2C-survived fate, while those inheriting more 53BP1 foci tended to adopt the 2C-death cell fate (85.19%, *p* = 0.008) (Figure 2I, column 3 to 4, 2K and Video S4). This asymmetric segregation led to an overall lower count of 53BP1 foci in the 2C-survived cells than in the 2C-death cells (Figure 2L). In addition to cells spontaneously entering the 2C-like state, DNA damage induced 2C-like cells also show asymmetric segregation of 53BP1 foci and the association of 2C-death fate with high 53BP1 foci number (Figures S3F-S3I). Altogether, these results demonstrate a strong correlation between asymmetric inheritance of DNA damage and the asymmetric cell fates observed through the 2C-like state.

How does DNA damage asymmetrically segregate into two sisters? We hypothesize that the asymmetric segregation relies on the sensing of DNA damage. By treating cells with inhibitors of ATM, ATR, CHEK or PARP, we observed that silencing the DNA damage response significantly reduced the fraction of 2C-like cells generating asymmetric daughter lineage fates (Figure 2M). Among the 2C-like cells that do give rise to asymmetric fates, the division of 53BP1 foci is less asymmetric in cells treated with ATM, ATR, CHEK or PARP inhibitor compared to control cells (Figure 2N). Together, these data suggest that DNA damage response is essential for asymmetric segregation of DNA damage, which in turn mediates the asymmetric fates of daughter lineages.

### Functional heterogeneity of 2C-like cells

To test the prediction of functional rejuvenation, we assessed the functional status of mESCs going through different stages of the 2C-like cycle. First, we examined whether 2C-death and 2C-survived fates can be distinguished by the fluorescent intensity of 2C reporters in a snapshot. We sorted cells with high 2C reporter intensity (reporter-high, top 0.5% for MERVL and top 1% for Zscan4) and lower, but above-background 2C reporter intensity (reporter-low, 98.5-99.5 percentile for MERVL or 98-99 percentile for Zscan4) into separate wells and followed the fates of these cells using live-cell imaging (Figures 3A-3B). More than 70% 2C reporter-high cells died within 48 hours, consistent with the 2C-death fate (Figure 3C). The rare survived reporter-high cells have fewer 53BP1 foci than the majority that died (Figure S3J). In contrast, 2C reporter-low cells showed comparable probability of dying, returning to a reporter negative state, or undergoing asymmetric division (Figure 3D). Therefore, while 2C-death cells can be readily identified in a snapshot by their high reporter intensity, cells with low reporter intensity comprise a mixture of those entering the 2C-like state that would undergo asymmetric division, 2C-death cells that are increasing their 2C reporter intensity and 2C-survived cells (Figure 3E).

**Figure 3.**
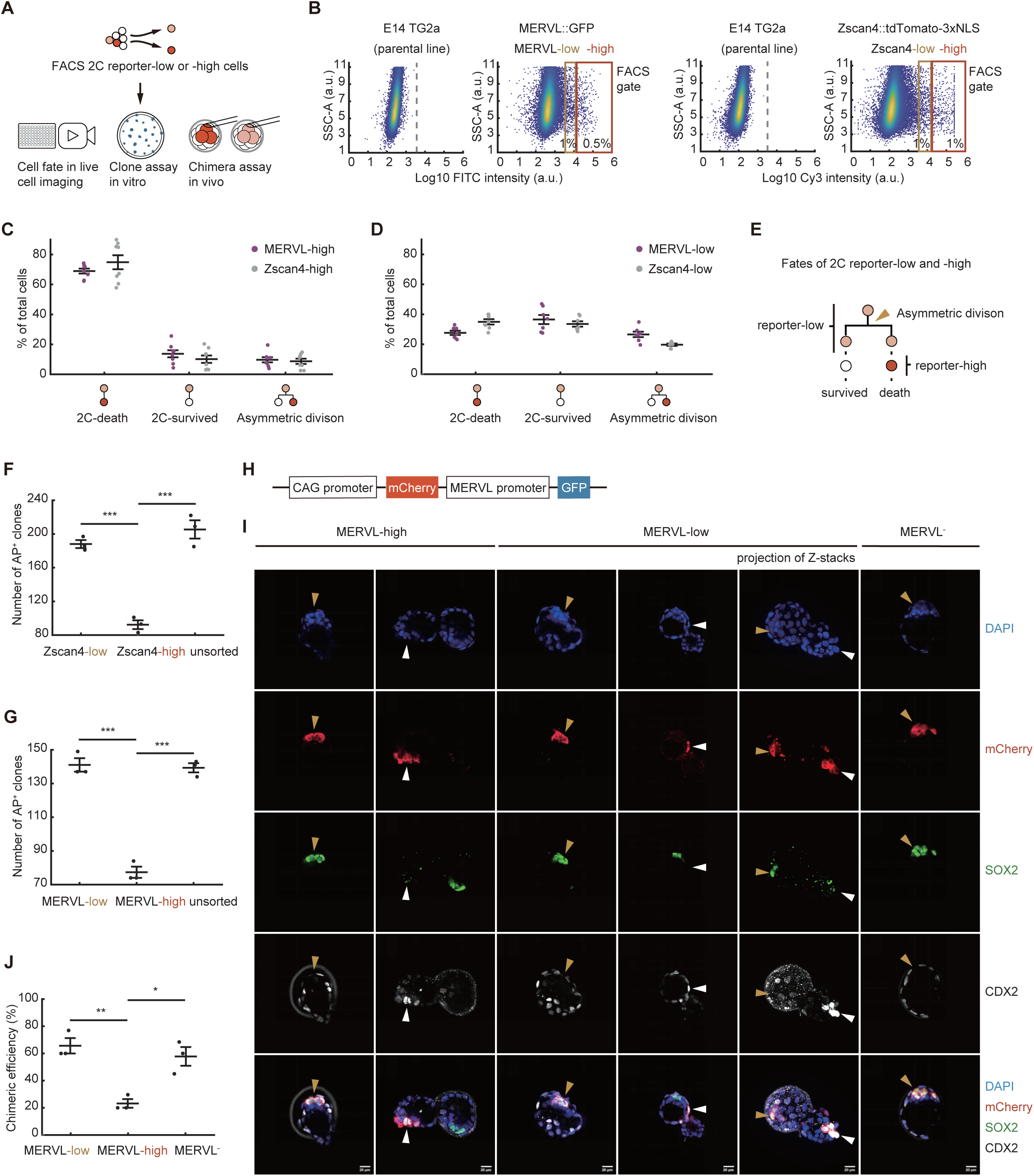
Functional heterogeneity of 2C reporter^+^ cells. (A) Experiment design of (B)-(J). 2C reporter-high or –low subpopulations were sorted by FACS for live-cell imaging, clone assays, and chimera assays. (B) Sorting gates for 2C reporter-low or –high subpopulation in MERVL::GFP (left) and Zscan4::tdTomato-3xNLS (right) cell lines. (C, D) Percentage of 2C-death, 2C-survived, and asymmetric division in 2C reporter-high (C) or –low (D) cells within 48 hours after FACS. (E) A schematic to depict the fates of 2C reporter-low and –high cells. The arrowhead indicates the asymmetric division. (F, G) Clone assays of 2C reporter-low cells, 2C reporter-high and unsorted cells (F, Zscan4 reporter; G, MERVL reporter). The numbers of AP^+^ colonies were quantified. (H) The construct used in experiments of (I)-(J). (I) Immunofluorescent staining of chimeric blastocysts incorporating donor cells (labeled by mCherry) from sorted MERVL-low, MERVL-high, and MERVL^−^ cells. SOX2, ICM marker; CDX2, TE marker. Yellow arrowheads indicate mCherry^+^ SOX2^+^ cells; white arrowheads indicate mCherry^+^ CDX2^+^ cells. Scale bar, 20 µm. Images are from single Z position with the exception of column 5 that is the maximum intensity projection of Z stacks. (J) Chimeric efficiency of MERVL-low, MERVL-high and MERVL^−^ cells. Error bars indicate mean ± SEM (C-D, F-G, J). Two-tailed unpaired student t-test (F-G, J). See also Figure S3.

Functionally, reporter-high cells generate substantially fewer Alkaline Phosphatase (AP) positive clones than unsorted and reporter-low cells in clone assays (Figures 3A and 3F-3G). To further test their ability to contribute to embryonic development, we injected them into 8-cell embryos and tested chimera formation at the blastocyst stage (Figure 3A). Both reporter-high and reporter-low cells are able to contribute to the trophectoderm (TE) and inner cell mass (ICM), in contrast to reporter negative cells that exclusively contribute to ICM (Figure 3I). However, compared to reporter-low cells, a significantly smaller fraction of reporter-high cells contributes to chimera formation (Figure 3J). These results suggest that cells with high 2C reporter intensity primarily adopt a dying fate and are less capable of contributing to embryonic development, supporting our model that they represent the lineage enriched with damage.

### Rejuvenation of 2C-exited cells

Next, we investigate the functional and molecular status of cells exiting from the 2C-like stage. Since cells with low 2C reporter intensity are a mixture of 2C-survived, 2C-entering and early stage of 2C-death cells, they cannot be separated based on the 2C reporter intensity alone. To sort 2C-survived cells and their recent descendants, we employed a tandem fluorescent protein timer, which encodes the temporal dynamics of transcription into the ratio of two fluorescence signals in a snapshot^26,27^. Specifically, this sensor consists of a fast-maturing and fast-degrading fluorescent protein (FP) – sfGFP-PEST, and a slow-maturing and slow-degrading FP – DsRed2, both expressed under the control of the MERVL promoter (Figure 4A). Upon entry into the 2C-like state, the MERVL promoter switches on, simultaneously producing both FPs, but sfGFP-PEST matures before DsRed2. Following deactivation of the MERVL promoter, DsRed2 persists within cells for much longer than sfGFP-PEST (Figure 4B). Therefore, the fluorescence ratio between the two FPs reflects the time relative to the on/off status of the MERVL promoter (Figures 4C and S4A).

**Figure 4.**
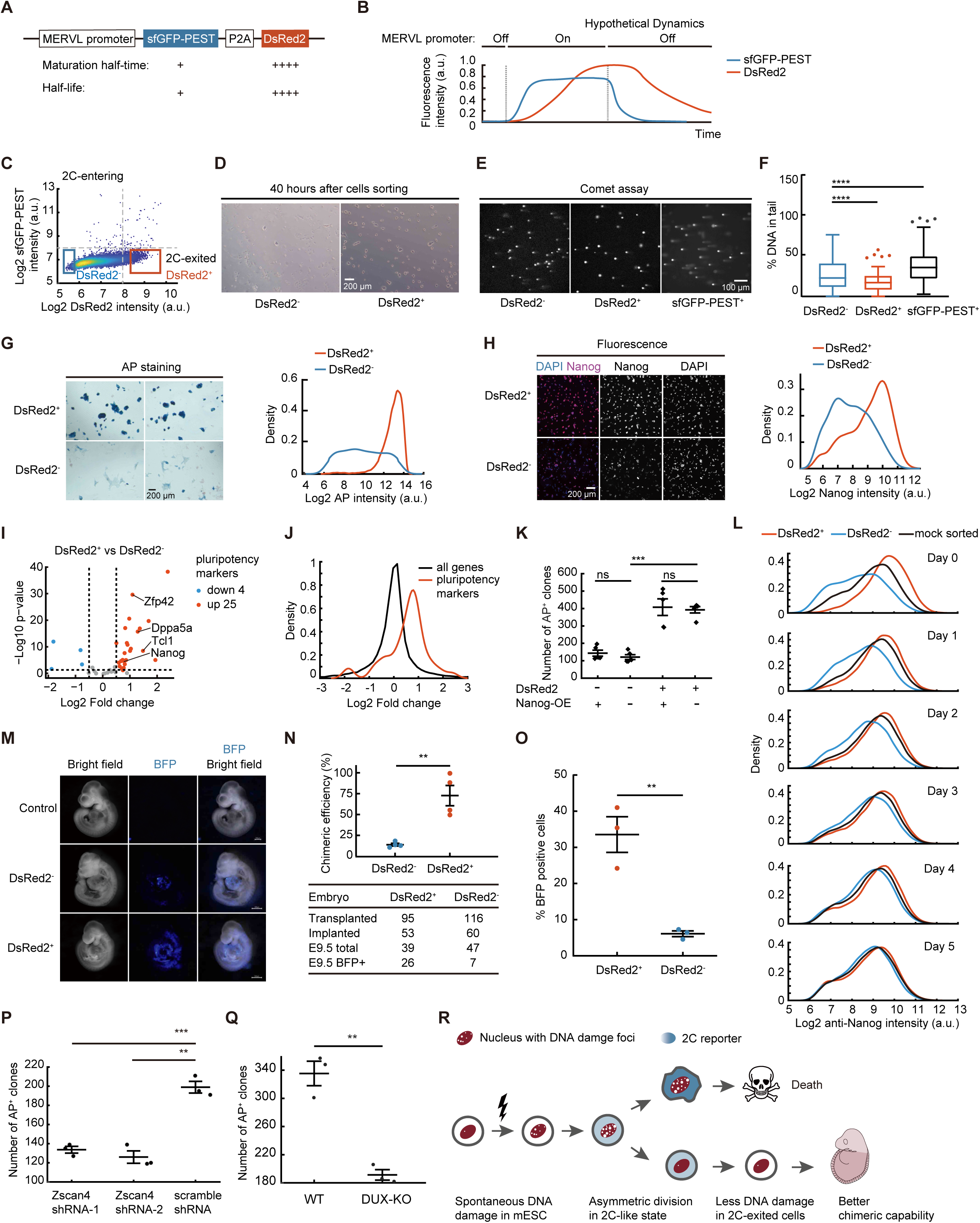
Cells recently exited the 2C-like state show improved pluripotency. (A) Design of the fluorescent timer reporter. (B) Schematics to show the hypothetical temporal dynamics of the two fluorescent proteins when cells pass through the 2C-like state. (C) Density scatter plot of sfGFP-PEST versus DsRed2 intensity in single cells. The red and blue box indicate the sorting gate for the DsRed2^+^ sfGFP-PEST^−^ and DsRed2^−^ sfGFP-PEST^−^ subpopulation (4% of total for each), respectively. (D) Clone morphology of sorted subpopulations in (C) cultured in standard conditions for 40 hours after FACS. Scale bar, 200 µm. (E, F) Comet assay of sorted subpopulations in (C); representative images (E) and quantification of the tail DNA fraction (F). n = 159 for each group. Scale bar, 100 µm. (G) AP staining of sorted subpopulations in (C). Left, representative images in brightfield; right, density distribution of AP intensity in Cy5 fluorescent channel. n > 4,000 for each group. Scale bar, 200 µm. (H) Nanog antibody staining of sorted subpopulations in (C). Left, representative images; right, quantification. n > 6,000 for each group. Scale bar, 200 µm. (I) Volcano plot of pluripotency markers in subpopulations sorted in (C). (J) Density distribution of the fold change of pluripotency marker genes and all genes. (K) Clone assays of sorted subpopulations in (C) with cells expressing tetracycline inducible Nanog. The numbers of AP^+^ colonies are quantified. (L) Density distribution of Nanog antibody staining in 5 days after sorting and culturing subpopulations in (C). n > 10,000 for each group. (M) Representative images of non-chimeric and chimeric embryos incorporating donor cells (labeled by BFP) from sorted subpopulations in (C). Scale bar, 500 µm. (N) Chimeric efficiency of the two sorted subpopulations in (C). (O) Percentage of the BFP positive cells in chimeric embryos, quantified by FACS. (P) Clone assays of cells expressing Zscan4 shRNA or scramble control. The number of AP^+^ colonies are quantified. (Q) Clone assays of DUX knock-out (DUX-KO) and wild-type (WT) C57 mES cell lines. The number of AP^+^ colonies are quantified. (R) A model of asymmetric division in a 2C-like state rejuvenating mESC lineage. During routine proliferation, mESCs accumulate DNA damage over time. Upon entering a 2C-like state, they undergo asymmetric division that segregates damage asymmetrically. One daughter lineage becomes damage-enriched and undergoes cell death, while the other is functionally rejuvenated with reduced damage burden. Error bars indicate mean ± SEM (K, N-Q). Two-tailed unpaired student t-test (K, N-Q); Mann–Whitney U test (F). See also Figure S4.

Using this reporter, we sorted cells that had recently exited from the 2C-survived branch (DsRed2^+^ sfGFP-PEST^−^, Figure 4C). Compared to the subpopulation that had not recently passed through the 2C-like state (DsRed2^−^ sfGFP-PEST^−^), sorted 2C-exited cells showed better dome-shaped colony morphology and fewer signs of DNA damage (Figures 4D-4F). They also show more robust staining of pluripotency markers such as AP, Nanog and Oct4 (Figures 4G-4H and S4B-S4C). At the transcriptome level, pluripotent marker genes are overall upregulated in DsRed2^+^ cells compared to DsRed2^−^ cells (Figures 4I-4J). DsRed2^+^ cells also generate more AP^+^ clones in clone assays (Figure 4K). However, overexpression of Nanog does not rescue the reduced clone-forming capacity in DsRed2^−^cells (Figure 4K), suggesting that decreased Nanog expression is likely a consequence rather than a cause of the impaired pluripotency in DsRed2^−^ cells. To determine the longevity of the enhanced pluripotency in DsRed2^+^ cells, we sorted DsRed2^+^ and DsRed2^−^, and remeasured the distribution of Nanog intensity daily. Surprisingly, the two subpopulations took less than 5 days to re-establish the steady state of Nanog expression (Figure 4L), suggesting that the gained pluripotency has an average lifetime of fewer than 10 generations.

We further tested the pluripotency of 2C-exited cells in vivo in chimera assays. When injecting 2C-exited cells into mouse blastocysts, 73.3% embryos were chimera with contribution from the donor cells (BFP^+^), a much higher efficiency than 12.8% using the DsRed2^−^ cells (Figures 4M-4O, and S4D-S4G). Collectively, these results indicate that cells survive from the asymmetric divisions and recently exited the 2C-like state show enhanced pluripotency, akin to a functionally rejuvenated state.

It remains controversial whether passing through the 2C-like state is essential for maintaining long-term pluripotency of mESC culture^8,9^. We tested this by knocking down Zscan4 (Figures S4H-S4I) or knocking out DUX (Figure S4J) to suppress entry into the 2C-like state. Both perturbations greatly reduced MERVL^+^ cells^11^ (Figures S4K-S4L), and resulted in the attenuated pluripotency of mESCs as shown in clone assays (Figures 4P-4Q). Together, our data suggest that entering the 2C-like state indeed represents an important mechanism for rejuvenating the embryonic stem cell lineage despite individual cell aging (Figure 4R).

## Discussion

Our data support a model wherein seemingly symmetrically dividing mESCs leverage asymmetric divisions during a sporadic 2C-like state to extend their lineages. By selectively concentrating damage into one daughter cells, this division generates two progenies of distinct functional age: a compromised one enriched for damage and a rejuvenated one with enhanced pluripotency.

Consistent with this model, our data demonstrate that 2C-like cells are functionally heterogeneous, exhibiting significant variability in their chimeric capability. This finding challenges the conventional view of 2C-like cells as a single population and reconciles seeming contradicting findings in the field: while 2C-like cells exhibit elevated DNA damage and cell death – suggesting a compromised state – blocking their formation triggers a culture crisis in mESCs, indicating their essential role in self-renewal. Our model unifies these observations by proposing 2C-like state produces divergent lineages – one destined for elimination that accounts for the observed damage, and another that ensures lineage renewal.

This model represents a solution for managing damage in immortal cell lineages, and to our knowledge, the first such kind in mammalian cells. While asymmetric division is well established in morphologically asymmetric systems like budding yeast, it remains unknown whether other morphologically symmetrically dividing cells have the potential of generating daughter cells with distinct ages. The findings of asymmetric divisions in simple organisms, such as E. coli, imply that the prevalence of such division and its association with cellular aging might be more widespread than previously envisioned^28^. Although the 2C-like state may be unique to mESCs, we hypothesize that the downstream machinery mediating asymmetric damage segregation could be conserved across various immortal lineages. From bacteria aging to germline-soma segregation, and even the queen-worker differentiation in eusocial insects like bees, the asymmetric division in mESCs exemplifies a broader principle: progeny asymmetries at different levels can function to generate diversity for selection within a population, allowing lineage renewal at the expense of a disposables subset. As the first such example in mammalian cells, mESCs offers unique model for studying cellular aging and rejuvenation.

Historically, telomeres have been recognized as the primary determinant of replicative lifespan. However, the discovery of telomerase and alternative lengthening mechanisms suggests that telomere imposes a signaling limit rather than an absolute barrier^29,30^. Instead, spontaneous cellular damage is an inevitable source of cellular aging. While our data suggest that asymmetric divisions serve as a critical mechanism to partition this damage, its capacity may thus act as a key control knob of lineage renewal, with practical consequences for disease and regeneration. For example, targeting stem cell exhaustion – a hallmark of organismal aging and a major hurdle in regenerative medicine – our findings highlight a potential avenue by inducing asymmetric division in stem cells. Conversely, in cancer, blocking undesired lineage rejuvenation may be attainable by manipulating their asymmetric divisions. Intriguingly, the two-cell markers DUX4 and Zscan4 have been found to be expressed in many human cancers, often in a small subpopulation^31^. Whether asymmetric division occurs in this context is open for further study. The precise mechanism governing asymmetric division also warrants exploration. The 2C-like subpopulation of mES cells presents a valuable model to dissect the machinery and variables that contribute to the differential partitioning of damaged components, as well as the broader underlying mechanisms of maintaining lineage fitness.

## Acknowledgments

We thank Dr. Yangming Wang for the Zscan4 promoter plasmid, Dr Ian Chambers for the E14 TG2a cells, Drs. José Silva and Sabrina Spencer for insightful comments on the manuscript, Ms. Chan Feng for manually verifying ESC lineages from cell tracking data, and the Min Lab and all members of CCLA for general discussions and assistance. This work is supported by National Natural Science Foundation of China (32170732, 32300665), Pearl River Talent Recruitment Program (2021ZT09Y233) and Major Project of Guangzhou National Laboratory (GZNL2024A02005).

## Author contributions

Conceptualization: MZ, MM; Methodology: AG, JL, JC, GW; Investigation: XW, FH, QS, ZX, BH, XZ, CD, LP, MDZ, TP; Visualization: XW, FH, QS, ZX, XZ; Funding acquisition: AG, MZ, MM; Project administration: MZ, MM; Supervision: ZZ, MZ, MM; Writing – original draft: XW, HF, QS, ZX, MM; Writing – review & editing: XW, HF, QS, BH, JL, MZ, MM.

## STAR Methods

### RESOURCE AVAILABILITY

#### Lead contact

Further information and requests for resources and reagents should be directed to and will be fulfilled by the lead contact, Mingwei Min.

## Materials availability

All materials generated in this study are available from the lead contact under material transfer request.

## Data and code availability

- Numerical data from image analysis will be deposited in BioStudies database (http://www.ebi.ac.uk/biostudies) and publicly available as of the date of publication.
- Raw image data generated in this study will be shared upon request.
- Code for image processing is on Github (https://github.com/tianchengzhe/EllipTrack, https://github.com/scappell/Cell_tracking).
- Raw sequencing data are deposited to GEO database and will be available with accession number GSE287152 and GSE289323.
- Any additional information required to reanalyze the data reported in this study is available from the lead contact upon request.

## EXPERIMENTAL MODEL AND STUDY PARTICIPANT DETAILS

### Mice

The mice were fed a normal diet and housed with 20°C–25°C temperature, 30–70% humidity, and a 12-hour light-dark cycle. All procedures related to animals were performed following the ethical guidelines of the Guangzhou National Laboratory and approved by the Guangzhou National Laboratory Animal Care and Use Committee.

### Cell culture

Unless specified otherwise, mES cells were maintained in serum/LIF medium composed of high glucose DMEM supplemented with 15% fetal bovine serum (FBS), 1 mM sodium pyruvate, 0.1 mM non-essential amino acids (NEAA), 2 mM GlutaMAX, 0.1 mM β-mercaptoethanol and 25 ng/mL mouse leukemia inhibitory factor (mLIF). The serum-free 2i/LIF medium contained 1:1 mix DMEM/F12 and Neurobasal with 0.1 mM NEAA, 2 mM GlutaMAX, 0.5× N2 supplement, 0.5× B27 supplement, 25 ng/mL mLIF, 3 μM CHIR-99021, 1 μM PD032590, and 0.1 mM β-mercaptoethanol. The medium was changed daily. Cells were passaged with 0.05% Trypsin-EDTA every 2-3 days. Culture plates were coated with 0.1% gelatin for at least 20 minutes at 37°C. For live cell imaging, phenol-red free DMEM or DMEM/F12 was used to reduce background fluorescence. For C57 ES cell lines, 2i was added into the serum/LIF medium. All cells were cultured in a humidified incubator at 37°C and 5% CO_2_. Mycoplasma was routinely tested using Mycoplasma Detection Kit to ensure mycoplasma-free culture conditions.

## METHOD DETAILS

### Construction of stable cell lines

To establish MERVL::GFP, Zscan4::tdTomato-3×NLS, MERVL::sfGFP-PEST-P2A-DsRed2 reporter cell lines, we cloned the MERVL promoter or the Zscan4 promoter into PiggyBac plasmids, positioning them upstream of the respective fluorescent protein coding sequences. The MERVL promoter is as reported before^8^. The Zscan4 promoter covers between the *Zscan4d* start codon and the 2570 bp upstream site from the start codon. The PEST sequence is used for rapidly degrading sfGFP^32^. The MERVL::tdTomato plasmid was kindly shared by Dr. Samuel Pfaff on addgene (40281)^8^. The mCherry driven by CAG promoter was added to the MERVL::GFP construct for chimera assays in Figure 3. The resulting constructs were transfected into E14 TG2a ES cells using Lipo3000. We subsequently isolated single cells by fluorescence-activated cell sorting (FACS). Clones with moderate expression levels of the reporter were used in imaging experiments. For chimera assays of DsRed2^+/−^ cells in Figure 4, the PiggyBac plasmid containing fluorescent protein BFP under the control of EF1α promoter was transfected into the MERVL::sfGFP-PEST-P2A-DsRed2 reporter cell line to label donor cells. For cell tracking, H2B-mIFP^33^ was introduced into 2C-reporter cells using lentivirus transduction. Lentivirus was also used to transduce the PIP-FUCCI-mCherry^21^ into MERVL:: GFP cells to monitor the cell cycle phases, and the 53BP1-mVenus into MERVL::tdTomato cells to monitor DNA damage. The 53BP1-mVenus fragment is a tandem Tudor domain of 53BP1 (protein sequence from 1218 to 1715)^34^ fused to the mVenus fluorescent protein. To overexpress Nanog, Nanog-T2A-EGFP was cloned into PiggyBac plasmids with Tet-on induced expression system and transfected into MERVL::sfGFP-PEST-P2A-DsRed2 cells. Transfected cells were then selected in 200 μg/mL hygromycin and induced Nanog expression with 1 μg/mL doxycycline. shRNA sequence against Zscan4 are as reported before^5,35^. The shRNA was introduced into E14 TG2a ES cells using lentivirus transduction, expressed under the control of U6 promoter. Transduced cells were then selected in 1 μg/mL puromycin for 5 days. The DUX-KO C57 ES cells were from a single clone, isolated from DUX^−/−^ blastocyst and genotyped by PCR.

### Drug treatment

The concentration of drugs: DTT, Dithiothreitol, 2 mM; TM, Tunicamycin, 10 μM; BTZ, Bortezomib, 1 μM; 2-MeOE2, 2-Methoxyestradiol, 10 μM; NaCl, 200 mM; Sucrose, 400 mM; 10.67% PEG, Poly (ethylene glycol); Aph, Aphidicolin, 1.25 μM, 2.5 μM and 5 μM; HU, Hydroxyurea, 0.2 mM, 0.4 mM and 0.8 mM; MMC, Mitomycin C, 0.25 μg/mL, 0.5 μg/mL and 1 μg/mL. ATR inhibitor VE-821, 0.5 μM; ATM inhibitor KU-5593, 10 μM; CHEH1/2 inhibitor Debromohymenialdisine, 20 μM; PARP inhibitor PARP-1-IN-3, 1.5 μM;. Aph, HU, and MMC treatment were carried out for 8 hours followed by wash-off. Cells were then cultured in fresh culture medium for another 24 hours before fixation. DTT, TM, BTZ, 2-MeOE2, NaCl, Sucrose, and PEG were induced for the time shown in the figures.

### Live cell imaging

Cells were seeded on a glass bottom 96-well plate, coated with mouse or human laminin for 8-12 hours before the imaging. Live cell imaging was performed on a Nikon Ti2-E microscope with a Spectra X light engine (Lumencor) and appropriate filter sets (mIFP: ex 640/30 nm, em 720/60 nm; tdTomato, mCherry or DsRed2: ex 550/15 nm, em 595/33 nm; GFP or mVenus: ex 475/28 nm, em 519/26 nm). Either a 10× 0.45 NA objective or a 20× 0.75 NA objective was used. Images were taken by an ORCA-Flash 4.0 V3 CMOS camera (Hamamatsu) at a frequency of 15 or 20 minutes per frame. Light exposure time was: 50-100 ms for Cy5, 30-50 ms for FITC, 30-50 ms for Cy3, with 50% lamp intensity. Light exposure and imaging frequency are minimized in the range with sufficient time resolution for single cell tracking and signal-to-noise ratio for quantification, while avoiding significant photobleaching and photodamage. During the imaging, cells were kept within a dark chamber stably maintained at 37°C, 5% CO_2_, and 95% humidity.

### Immunofluorescence (IF) and fixed cell imaging

For monolayer cell immunostaining, cells were fixed with 4% PFA, permeabilized with 0.2% TritonX-100 in PBS, blocked with 5% Goat Serum in 0.1% Triton X-100, and incubated with the primary antibody diluted in blocking buffer overnight at 4°C. Antibodies used: Phospho-Histone H2A.X (γH2AX, Ser139) (D7T2V), 1: 200; PERK (phospho T980), 1: 400; Phospho-p53 (Ser15), 1: 200; MuERVL-Gag, 1: 10,000; Nanog (eBioMLC-51), 1: 250; Oct3/4, 1: 200, Zscan4, 1:15,000; Alexa Fluor 488/568/647 Secondary Antibodies, 1: 1,000. Most IF experiments are imaged using the same wide field microscope setup as in live cell imaging. A laser scanning confocal microscope (ZEISS, LSM 900) was used to confirm the co-localization of γH2AX and 53BP1-mVenus. AP staining was carried out using the Vector Blue AP Substrate Kit following the manufacturer’s instruction. The bright-field images were captured with a ZEISS Axio Vert.A1 Inverted Microscope.

### Decay kinetics of the pluripotency rejuvenation

MERVL::DsRed2^+^, and MERVL::DsRed2^−^ cells were counted and seeded immediately after being sorted by FACS. For the next 5 days, cells were fixed at the same time every day with 4% PFA. IF were performed on the fifth day.

### Image processing, cell tracking, and signal extraction

Raw images were corrected to extract quantitative information from the fluorescence intensity. Briefly, dark noise was subtracted from the raw image followed by correction of illumination bias. Dark noise was measured by averaging 10 snapshots with the light source off. Illumination bias of each fluorescence channel was deduced by averaging the cell-free regions of all images within the channel. The background was then removed using top hat filtering.

Cells are segmented using the pre-trained model in Stardist^36^, with the nuclear channel as the input image (H2B-mIFP in live cell imaging and DAPI in fixed cell imaging). Between frames where the imaging plate was removed from and put back in the microscope (for example, during drug addition, medium change or IF after live-cell imaging), the plate jitter was calculated by registering images of the nucleus-stained channel and corrected prior to tracking. Cell tracking was carried out using a local tracking method^37^, or EllipTrack^38^. Briefly, the local tracking method tracks cells by screening the nearest future neighbors. EllipTrack employs machine learning techniques to deduce probabilities of cell overlap, cell migration, and mitosis using information of cell morphology and position, and construct tracks that maximize the probability throughout the entire movie using the Viterbi algorithm. The result from this global tracking pipeline was further refined using a local correction module, through iteratively swapping every two cell tracks between every two neighboring frames and keeping those swaps that significantly increase the combined probabilities of all tracks. To ensure the accuracy of lineages, we manually verified and corrected all tracks quantified in this study.

Fluorescence intensity of each cell was then extracted from background-removed images as the median intensity of pixels within its nuclear mask (nuclear intensity) or a 4-pixel wide ring around the nuclear mask (cytoplasmic intensity). Nuclear intensity was used for downstream analysis when the fluorescence of interest locates within nuclei or throughout the entire cell, while cytoplasmic intensity was used when the targeted fluorescence locates exclusively in the cytoplasm.

### Single cell analysis

In time-lapse imaging, MERVL reporter negative cells are those that exhibit no fluorescent intensity above background levels. MERVL reporter positive cells are classified as 2C-entering, 2C-death and 2C-survived as followed: the 2C-entering cells refer to cells between the appearance of MERVL::FP and the asymmetric division; the 2C-death cells refer to cells between the asymmetric division and cell death, which often exhibit high levels of MERVL reporter intensity; the 2C-survived cells are the sister branch of the 2C-death cells, ranging from the asymmetric division to the disappearance of MERVL reporter fluorescence (Figure 1J).

In lineage survival analysis, if at least one offspring in a lineage survived, the lineage was considered to have survived; if all the offspring in a lineage died, the lineage was considered to have died.

To track the fates of 2C-like cells and visualize the lineage tree of MERVL::GFP^+^ cells, we tracked the MERVL::GFP dynamics for each cell. Robust lowess smoothing was applied on MERVL::GFP intensity with a window of 2.5 hours to reduce noise. The lineage trees were plotted using in-house MATLAB scripts.

To quantify the MERVL::GFP intensity among different subpopulations of mES cells, we extracted the MERVL::GFP intensity for each cell at each frame and calculated the median MERVL::GFP intensity over a cell cycle. For MERVL::GFP^+^ cells, we included all cell cycles within each dying or alive lineage from the point of asymmetric division to cell death (2C-death lineage) or returning to MERVL::GFP^−^ state (2C-survived lineage).

PIP-FUCCI and H2B reporters were used to determine the boundary of cell cycle phases. The M/G1 boundary was recognized by chromosome segregation, indicated by the H2B signal. The G1/S boundary was recognized by the fall of PIP-FUCCI signal below the background threshold. The S/G2 boundary transition was defined as the time PIP-FUCCI level begins to rise, detected using a triangle thresholding method^39^.

To calculate the percentage of MERVL::GFP^+^ cells for each frame in exogenous cellular damage movie, the threshold for defining MERVL::GFP^+^ is set as followed: *T = μ + k*σ*, where the *μ* is the median of all cells in the control condition and *σ* represents its standard deviation. The *k* is a constant with a value of 2.7.

To track 53BP1 foci numbers in 2C asymmetric division lineages, 2C lineage fates were manually verified and 53BP1 foci numbers were counted manually in the S phase.

In all box plots, center line indicates median; box limits indicate upper and lower quartiles; whiskers show 1.5x interquartile range; points show outliers.

Numerical data analysis was carried out in MATLAB and plotted in MATLAB, Origin or GraphPad.

### Comet assay

The different subpopulations of 2C-like cells were sorted using FACS. Comet assay was carried out using a Comet Assay Kit, following the manufacturer’s instruction. Briefly, cells suspension was prepared at the density of 1,000 cells/μL, mixed with low-melting agarose in a volume ratio of 1: 7.5 and then put in solid normal agarose at 4°C for 10 minutes until agarose becoming solid. Cells were lysed in cold lysis buffer overnight. DNA was unwound in running buffer (200 mM NaOH and 1 mM EDTA) for 1 hour at room temperature, followed by electrophoresed at 20 V within cold running buffer for 30 minutes, and neutralized with neutralization buffer (0.4 M Tris-HCl, PH 7.5) for 5 minutes. The slides were imaged on a Nikon Ti2-E microscope after DAPI staining for 5 minutes in dark. Comet assay analysis was carried out using the OpenComet plugin^40^ in ImageJ^41^.

### Single cell RNA sequencing

MERVL::GFP^+^ and MERVL::GFP^−^ cells were sorted followed by construction of single cell RNA sequencing library using the Chromium Next GEM Chip G Single Cell Kit following the manufacturer’s instruction. The libraries were sequenced on an Illumina HiSeqXTen instrument using 150 nt paired-end sequencing. Overall sequence quality was examined using FastQC (v0.11.2) and adaptor sequences were clipped using Trim Galore (v0.6.4). The reads were mapped to the mouse genome (mm10) and a count matrix was generated using STAR (v2.7.6a)^42^. Low quality cells were filtered out based on a gene number threshold of 3000. Dimensionality reduction was carried out on the count matrix using Principal Component Analysis (PCA) followed by clustering and visualization using Uniform Manifold Approximation and Projection (UMAP). Differentially expressed gene analysis was performed using the Python package Scanpy (v1.8.1) with the function ‘rank_genes_groups’^43^. Genes showing more than two-fold expression change with an adjusted *p*-value < 0.05 were considered as differentially expressed. The list of p53 transcriptional target genes was compiled from a previous global run-on sequencing (GRO-Seq) study in human cells^44^ and mapped to their mouse homologs using biomaRt (v2.50.1)^45^. The volcano plots were generated using the seaborn package (v0.11.0)^46^ and labelled UMAPs using the Scanpy package. These marker genes sequences reference from NCBI (Zscan4c ID: 245109; Ddit4l ID: 73284; Nanog ID: 71950; SOX2 ID: 20674; Pou5f1 ID: 18999).

### Bulk RNA sequencing

MERVL::DsRed2^+^ and MERVL::DsRed2^−^ cells were sorted by FACS and lysed in Tri Reagent for RNA extraction. Three biological replicates were collected on different dates. Library construction and sequencing were done by China National Gene Bank. The library was constructed based on DNBSEQ Platform. In brief, after sample quality control, the mRNA was hybridized with oligo (dT) probe and captured by magnetic beads. The mRNA was then reverse-transcribed to first-strand DNA, which was used as a template to synthesize second-strand DNA. The dTTP-tailed adaptor was ligated to both ends of the dsDNA fragments. The ligation product was amplified by PCR, and circularized to get single-stranded circular (ssCir) library, which was then amplified through rolling circle amplification to obtain DNA nanoball (DNB). The DNB was then loaded to flowcell, and sequenced by DNBSEQ Platform. PE150 sequencing was used. Sequence quality was examined using FastQC (v0.12.1) and low-quality bases were removed using Trimmomatic (v0.39). The reads were aligned to mouse genome (GRCM39) using STAR (v2.7.11b). The alignment will be called if its ratio of mismatches to mapped length is less than or equal to 0.04. Mapped fragments were counted using RSEM (v1.3.3). Differential expression analysis was carried out using DESeq2 (v1.45.3) in R studio (v4.4.1). Genes with log2 (Fold Change) > 0.5 and adjusted *p*-value < 0.05 were considered as differentially expressed between the two cell populations. The list of pluripotent genes was from Peng Du et al^47^. The volcano plots were generated using the package tinyarray (v2.4.2).

### Clone assay

Cells were plated in 6-well cell culture plates at a density of 600 cells per well. The cells were cultured under our routine culture conditions for 5 days with medium change every two days. On the 5th day, the cells were fixed with 4% PFA followed by AP staining. Number of AP^+^ clones were counted manually in Fiji.

### Chimera assay

Six– to eight-week-old female mice were superovulated by injecting of 5-6 IU of pregnant mare serum gonadotrophin (PMSG). 46-48 hours after PMSG injection, 5-6 IU of human chorionic gonadotrophin (HCG) was administered. Superovulated female mice were set up for mating with ICR male mice. About 16-18 hours later, mouse zygotes were isolated in M2 medium and cumulus cells were removed with hyaluronidase. All of the embryos were then cultured in KSOM medium at 37°C under 5% CO_2_ in air under mineral oil. Approximately 6-8 FACS-selected DsRed2^−^ or DsRed2^+^ mES cells were injected into each E3.5 blastocyst. Injected embryos were then rest in KSOM medium. The next day, around 30 injected E4.5 embryos were transferred into both oviducts of an D0.5 pseudopregnant ICR mouse. Dissection was performed at E9.5. Chimeric conceptuses were observed using a Leica M165 FC Fluorescent Stereo Microscope and the manual scoring of chimeric embryos was blinded. For FACS quantification, cells were dissociated using a mixture of 0.05% trypsin:TrypLE (1:1) and neutralized in MEF medium (high glucose DMEM, 4 mM L-glutamine, 10% fetal bovine serum, 1 U/mL penicillin, 1 mg/mL streptomycin). The cells were subsequently centrifuged, and the pellets were resuspended in 2% knockout serum in PBS. After filtration through a 35 µm strainer cap, the cells were analyzed on a BD LSR Fortessa X-20. Data were analyzed using FlowJo 10 software.

### Blastocyst immunostaining

Approximately three FACS-selected MERVL::GFP^−^, MERVL::GFP-low or MERVL::GFP-high mESCs were injected into each E2.5 blastocyst. The injected embryos were then cultured in KSOM medium. Three days later, E5.5 embryos were fixed with 4% formaldehyde for 30 minutes at room temperature. Then, samples were washed three times for 5 minutes each with 0.1% Triton-X100 in PBS. After permeabilization with 0.3% Triton-X100 for 30 minutes, the samples were blocked with 3% donkey serum, 1% BSA and 0.1% Triton-X100 in PBS for 2 hours, and then incubated with the primary antibodies at 4°C overnight. The next day, embryos were washed four times with 0.1% Triton-X100 for 5 minutes each, incubated with fluorescent-labelled secondary antibodies for 2 hours and stained with DAPI for 5 minutes at room temperature. Antibodies used: CDX2 (D7T2V)/SOX2 (Btjce)/mCherry, 1: 200; Alexa Fluor 488/568/647 Secondary Antibodies, 1: 500. Images were taken using a Carl Zeiss LSM800 confocal microscope and processed using ZEN software. The chimeric efficiency is calculated as the percentage of mCherry^+^ blastocyst.

### Karyotype analysis

Karyotype analysis was performed by the karyotyping facility in Guangzhou Institutes of Biomedicine and Health. Briefly, mES cells were arrested at prometaphase with 200 ng/mL colchicine for 1 hour at 37°C, and collected in pre-warmed hypotonic KCl solution. After 30 minutes incubation at 37°C, cells were fixed with methanol and acetic acid (3: 1) and dropped onto pre-cooled slides. Chromosomes were stained with Giemsa solution, imaged using a ZEISS Axio Imager 2 Microscope and analyzed with the Ikaros Karyotyping Software. A minimum of 20 prometaphase spreads were analyzed.

## QUANTIFICATION AND STATISTICAL ANALYSIS

Quantification of imaging data was performed in MATLAB or ImageJ as described in the above methods section. Processed data are presented as the means ± standard error of the mean (SEM), median ± quantile (box plot) or single cell density distributions (density plot) as indicated in figure legends. Sample size and replicate numbers are indicated in figure legends. Statistical analysis is indicated in figure legends and main text. Legends of significance analysis: **** *p* ≤ 0.0001, *** *p* ≤ 0.001, ** *p* ≤ 0.01, * *p* ≤ 0.05, ns *p* > 0.05.

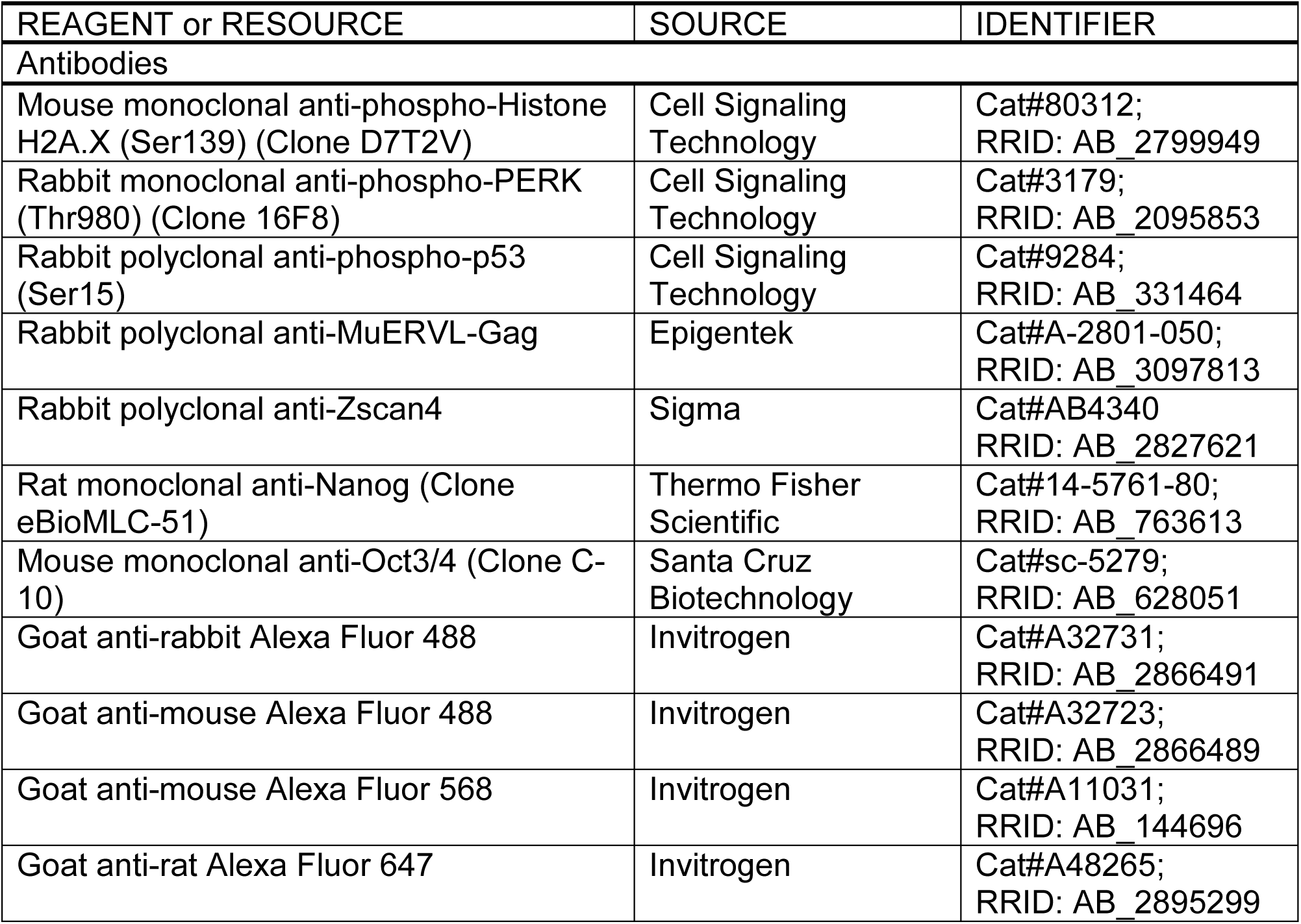
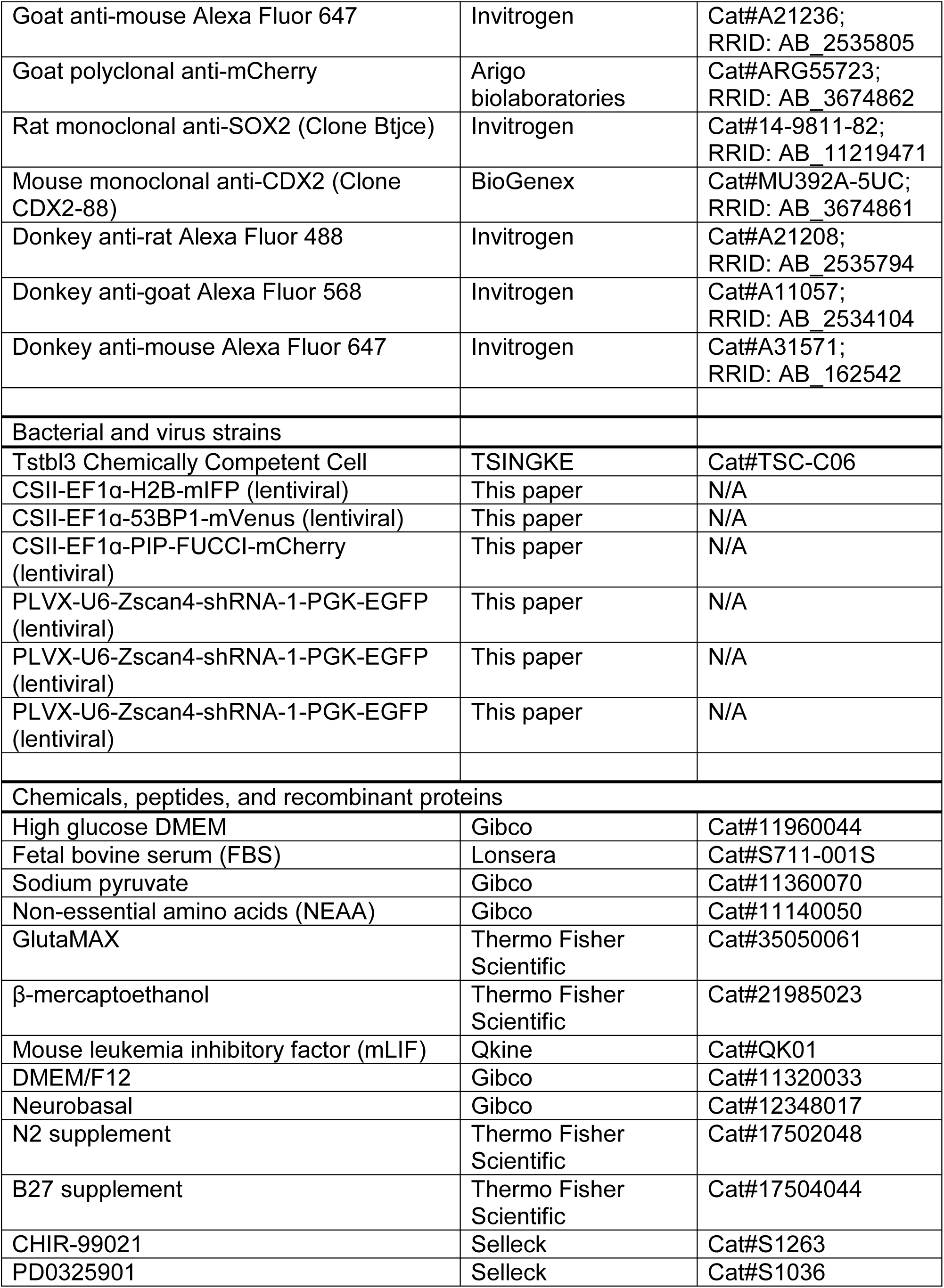
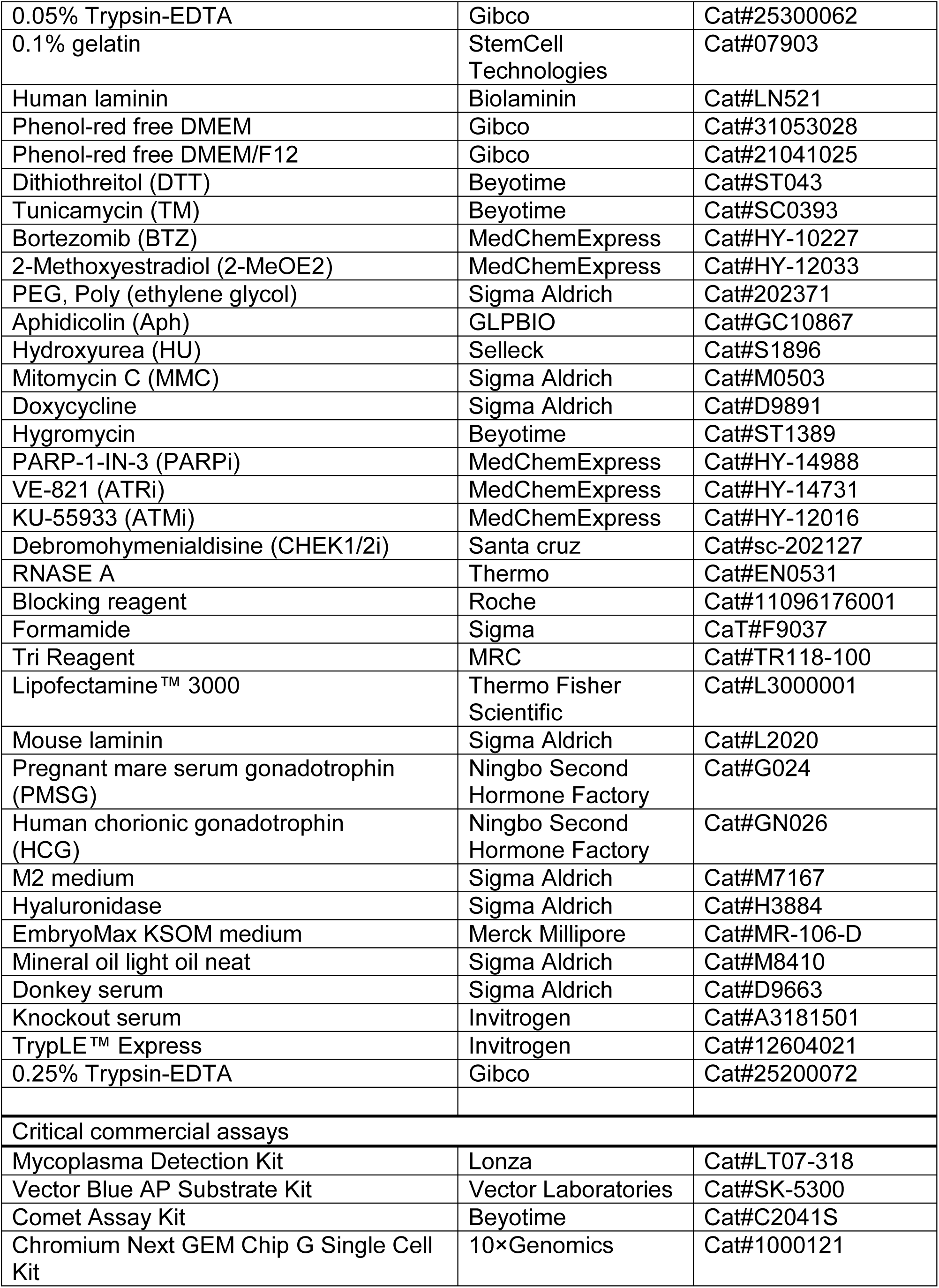
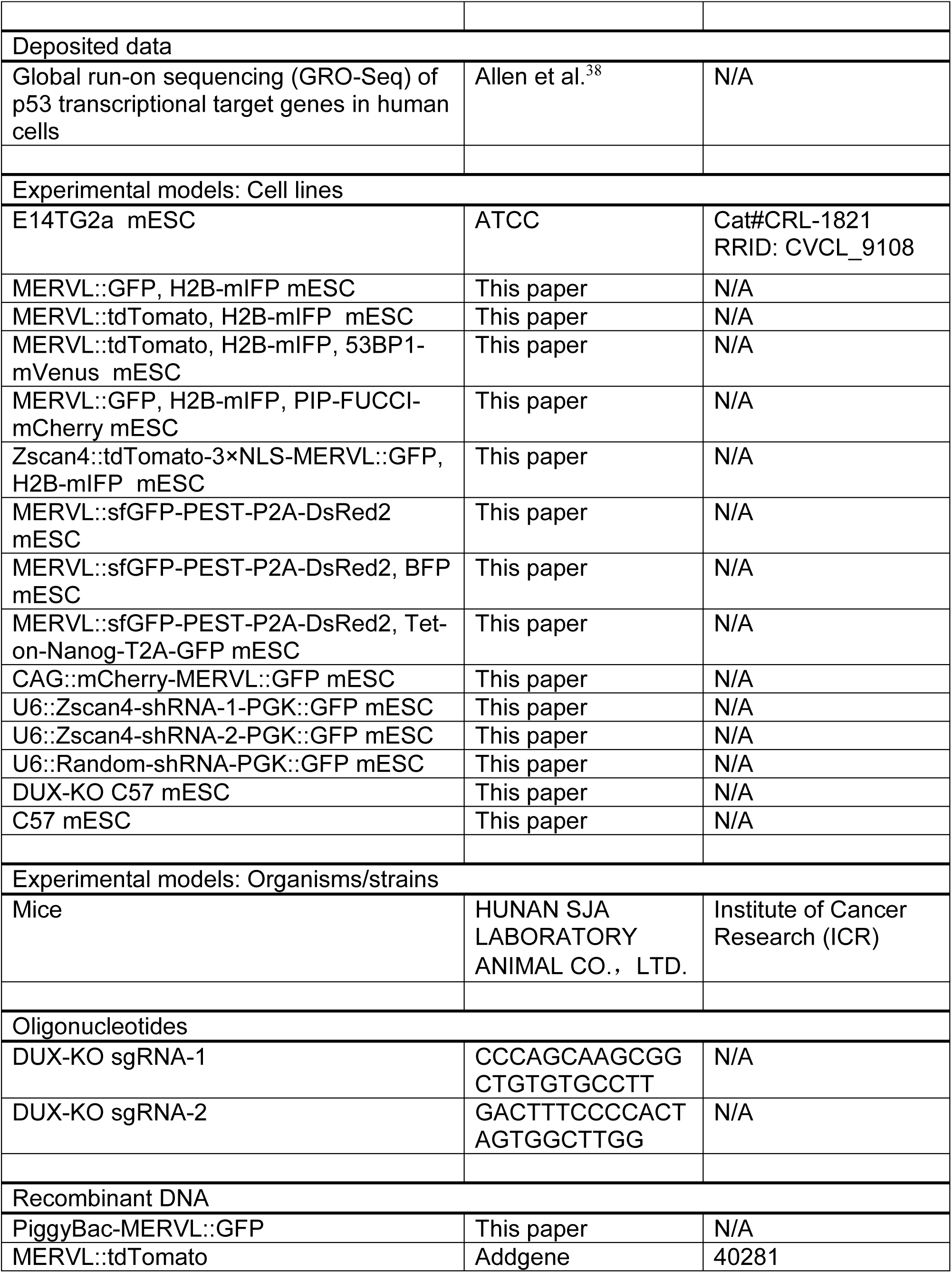
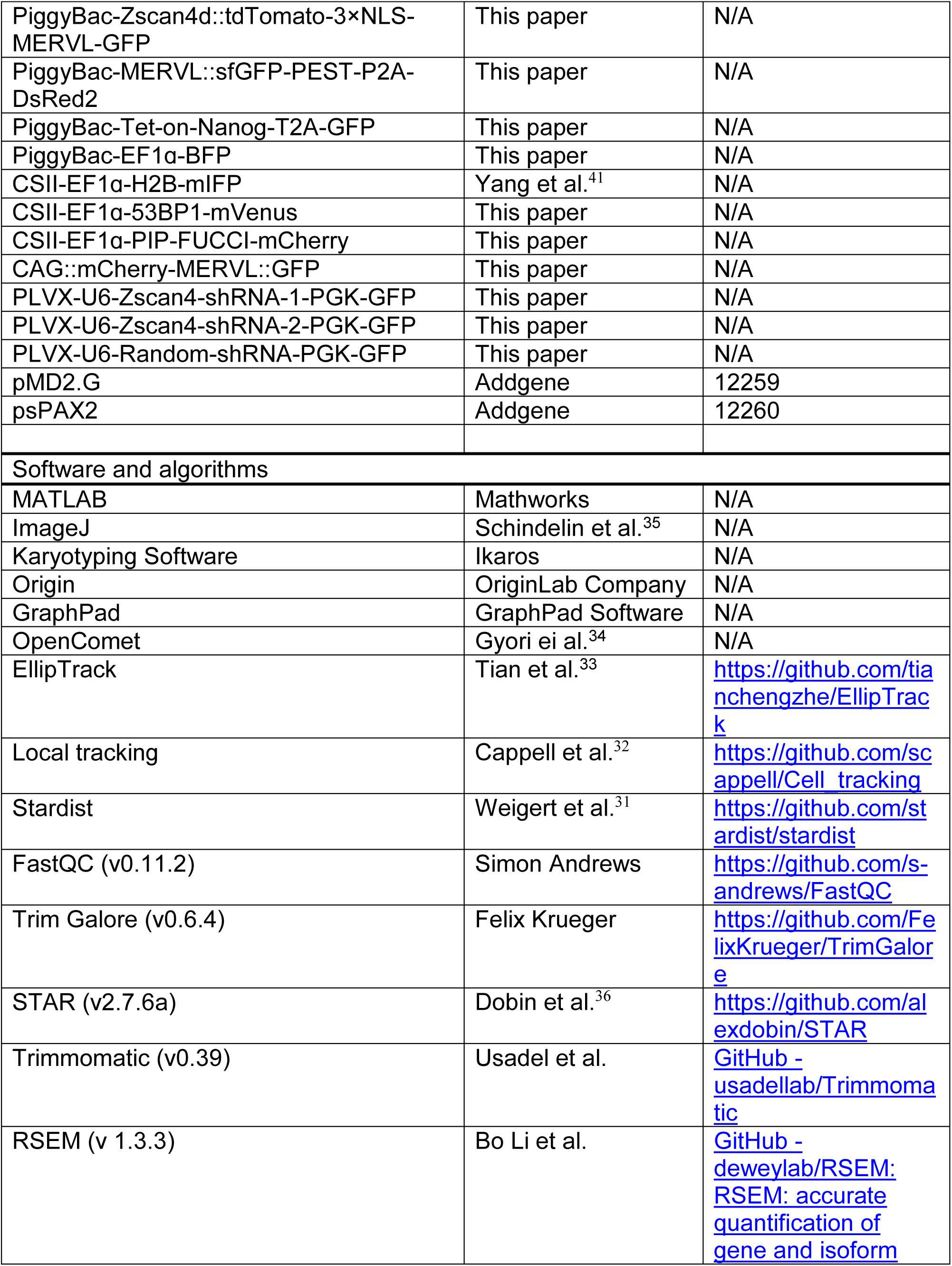
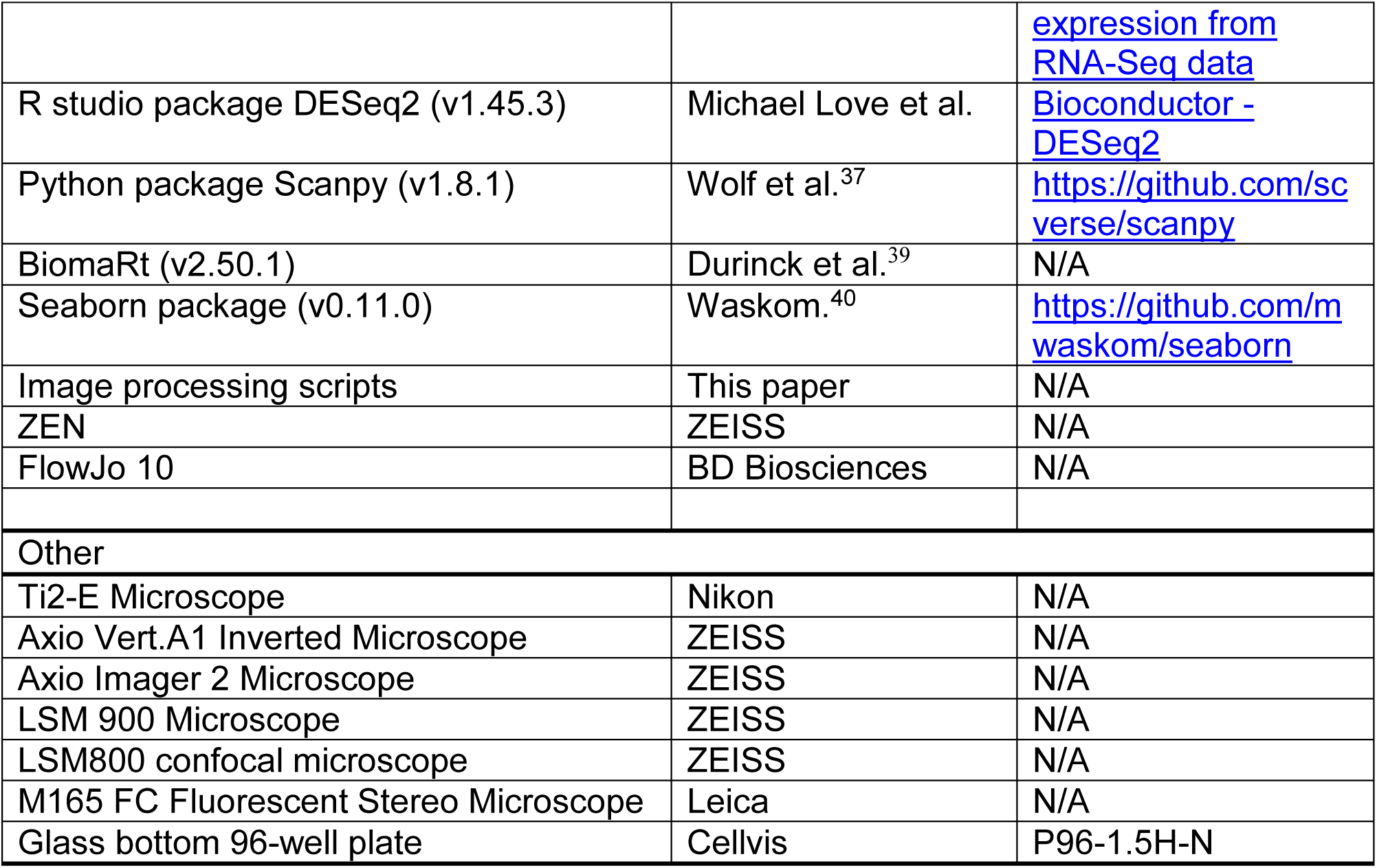
KEY RESOURCES TABLE.

### Supplemental information

#### Figures S1-S4

Figure S1. Additional examples of asymmetric daughter-cell fates in 2C reporter^+^ cells, related to Figure 1

Figure S2. Cellular damage in 2C reporter^+^ cells, related to Figure 2

Figure S3. Additional data on asymmetric segregation of DNA damage, related to Figure 2 and 3

Figure S4. 2C-exited cells show enhanced pluripotency, related to Figure 4

#### Videos S1-S4

Video S1. MERVL::GFP^+^ cells undergo cell death, related to Figure 1

Video S2. Distinct cell fates of MERVL^+^ sister cells, related to Figure 1

Video S3. Asymmetric division of Zscan4^+^ cells, related to Figure 1

Video S4. Asymmetric segregation of 53BP1 foci in MERVL^+^ cells, related to Figure 2

## Figure legends

**Figure S1.**
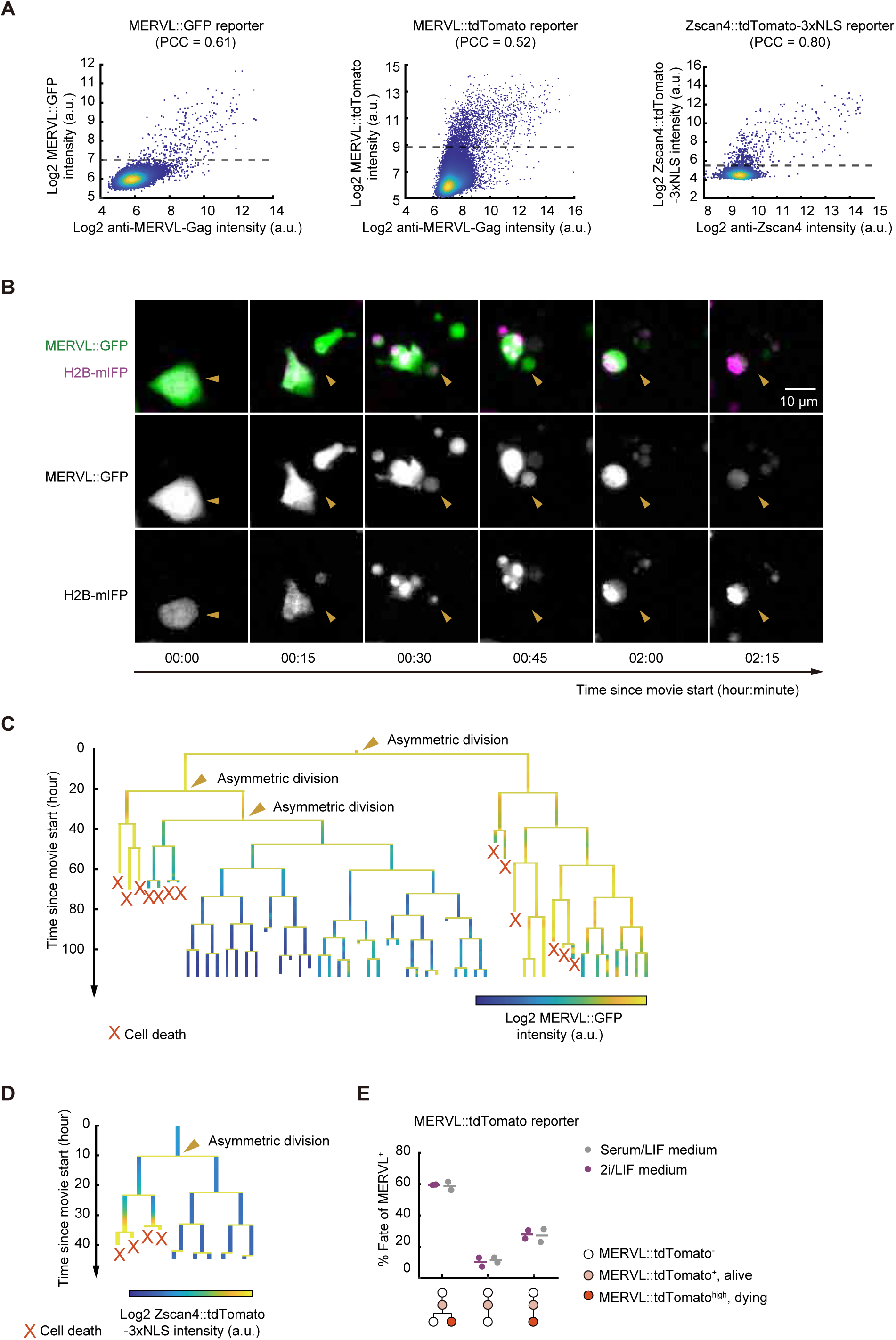
Additional examples of asymmetric daughter-cell fates in 2C reporter^+^ cells, related to Figure 1. (A) Density scatter plots of intensity from MERVL-Gag antibody staining versus MERVL::GFP (left), MERVL-Gag antibody staining versus MERVL::tdTomato (middle), and Zscan4 antibody staining versus Zscan4::tdTomato-3xNLS (right). PCC, Pearson Correlation Coefficient in reporter^+^ cells (above the dash line). (B) Enlarged images of cell death in MERVL::GFP^+^ cells. Cells undergo DNA condensation and fragmentation, with GFP intensity dropping to near background level between two frames (15 minutes). Scale bar, 10 μm. (C) An example lineage with consecutive asymmetric divisions in MERVL::GFP^+^ cells. The crosses mark cells that died; the arrowheads indicate asymmetric divisions. (D) An example lineage with asymmetric daughter-cell fates of Zscan4::tdTomato^+^ cells. The crosses mark cells that died; the arrowhead indicates asymmetric division; the heat color represents the MERVL::GFP intensity. (E) Asymmetric divisions also occur in mESCs cultured in the 2i/LIF condition. Each point represents a biological replicate. The lines indicate the mean.

**Figure S2.**
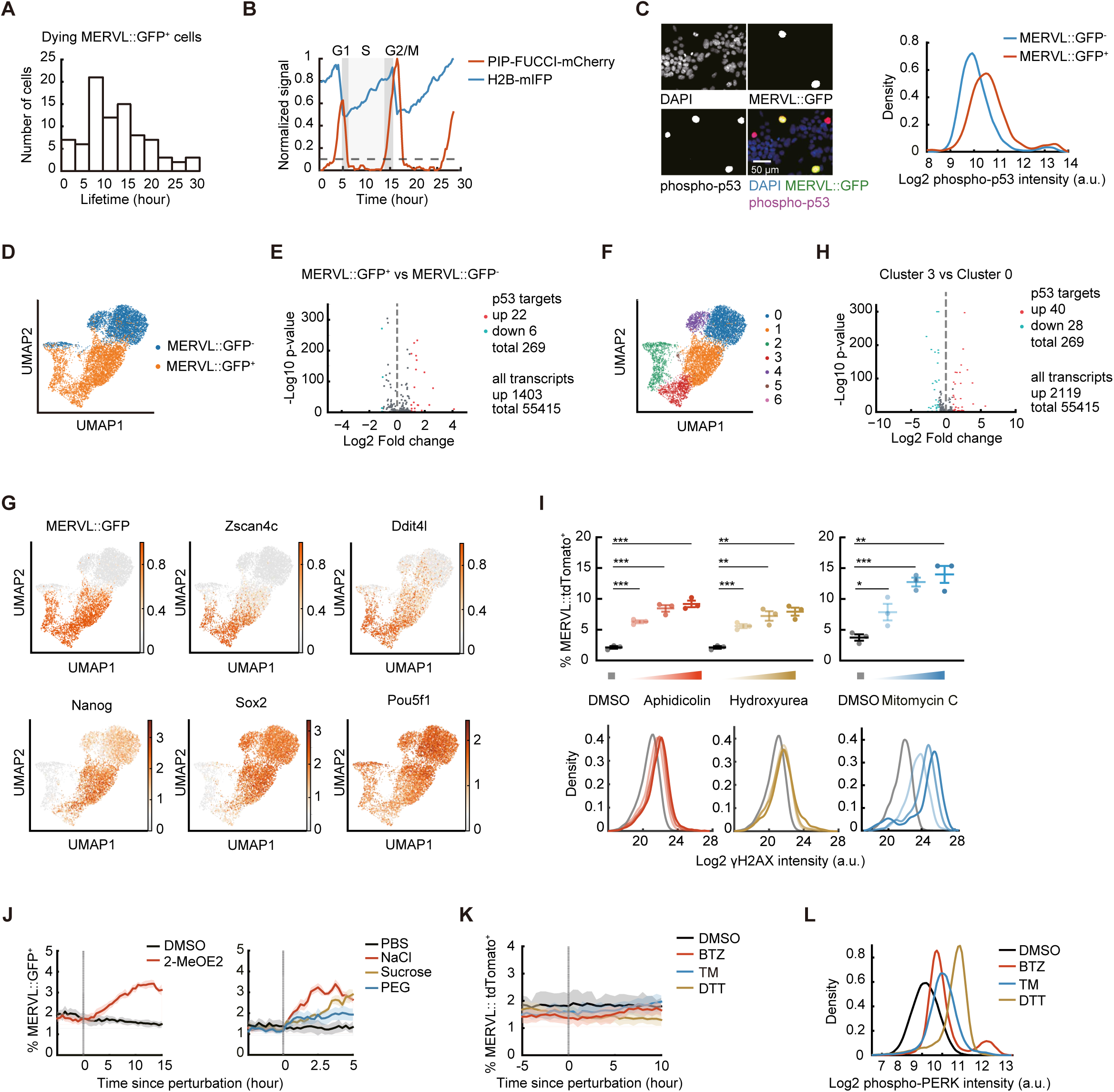
Cellular damage in 2C reporter^+^ cells, related to Figure 2. (A) The length of the last cell cycle in the dying MERVL::GFP^+^ cells, measured as the time between the last mitosis to cell death (n = 84). (B) The cell cycle dynamics of the PIP-FUCCI and H2B reporter. The drop in PIP-FUCCI reporter intensity marks G1/S transitions and its rise marks S/G2 transitions. The drop in H2B reporter intensity marks chromosome segregations. (C) Phospho-p53 staining in MERVL::GFP^+^ and MERVL::GFP^−^ cells. Left, representative images; right, density distribution of phospho-p53 intensity in single cell (610 MERVL^+^ cells and 60,342 MERVL^−^ cells). *p* < 0.0001. Scale bar, 50 μm. (D) Single cell transcriptomic profiling of sorted MERVL::GFP^−^ and MERVL::GFP^+^ cells. UMAP visualization shows distinct transcriptional states. (E) Volcano plot of p53 targets in MERVL::GFP^+^ versus MERVL::GFP^−^ cells. (F-G) A cluster (cluster 3) of MERVL::GFP^+^ cells expressing high levels of 2C-like markers, likely corresponding to the 2C-death cells. (H) Volcano plot of p53 targets in cluster 3 versus cluster 0 cells. (I) Top: the percentage of MERVL::tdTomato^+^ cells in DNA damage conditions, n = 3; bottom, the density distributions of mean γH2AX intensity in DNA damage conditions (n > 1,000 for each curve). (J) The percentage of MERVL::GFP^+^ cells upon applying oxidative (left) and osmotic stress (right), n = 4 for each group. (K) Inducing endoplasmic reticulum stress does not increase MERVL::tdTomato^+^ fraction over time (n = 4). BTZ, bortezomib; DTT, Dithiothreitol; TM, Tunicamycin. (L) The phosphorylation of PERK at T980 is induced by endoplasmic reticulum stress treatment in (K) (n > 9,000 for each condition). Error bars indicate mean ± SEM (I). Data are presented as 95% confidence intervals (J-K). Kolmogorov-Smirnov test (C); Two-tailed unpaired student t-test (I).

**Figure S3.**
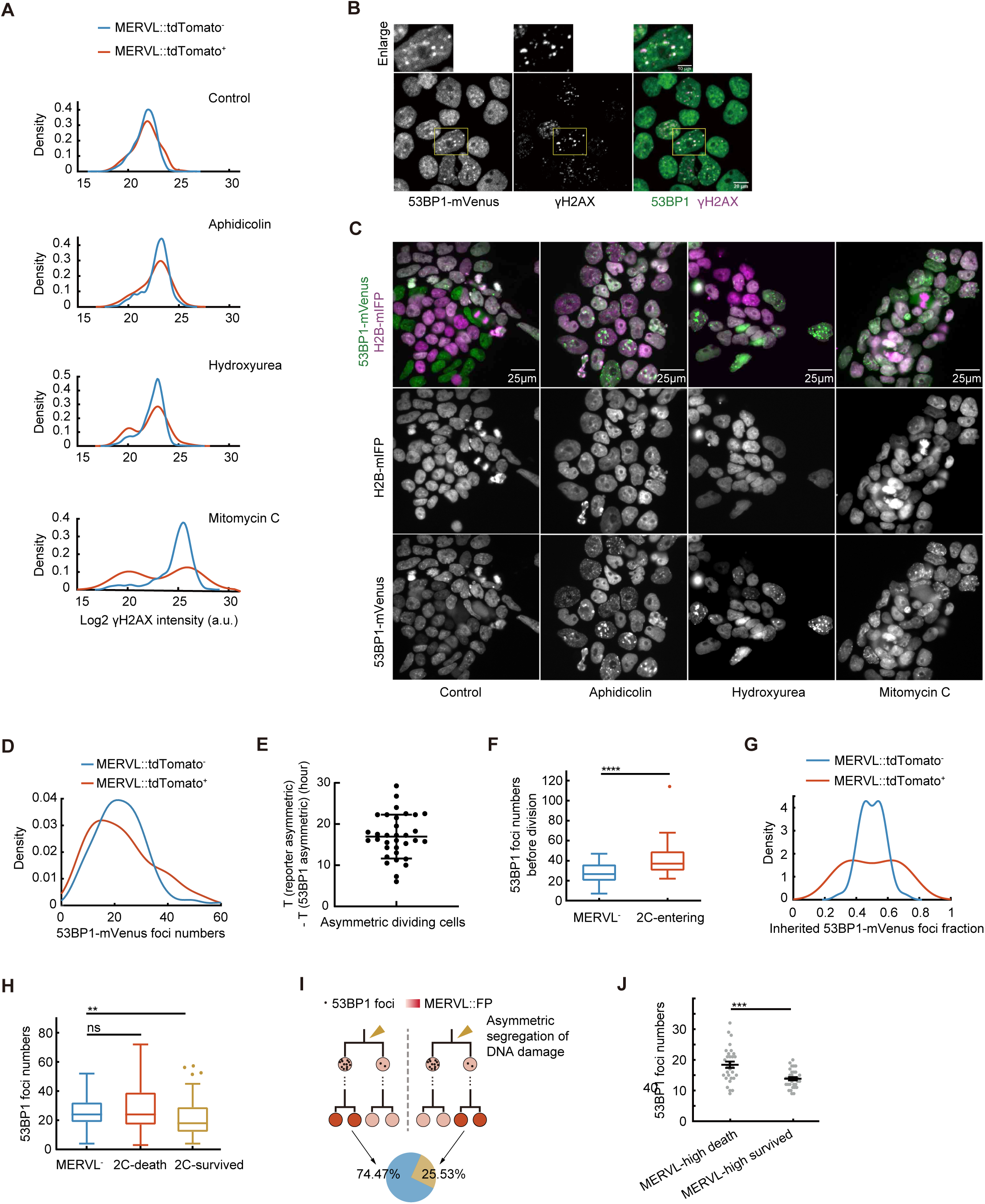
Additional data on asymmetric segregation of DNA damage, related to Figure 2 and Figure 3. (A) Density distribution of total γH2AX intensity in single MERVL::tdTomato^−^ and MERVL::tdTomato^+^ cells in control (n = 6,533) and induced DNA damage conditions: 6.4 μM Aphidicolin (n = 2,237), 0.32 mM Hydroxyurea (n = 2,306) and 1 μg/mL Mitomycin C (n = 645). (B) Co-localization of 53BP1-mVenus and γH2AX antibody staining. Scale bar of bottom row, 20 μm. Scale bar of enlarged images, 10 μm. (C) The number of 53BP1-mVenus foci increases upon treatment with DNA damage drugs (Aphidicolin, 6.4 μM; Hydroxyurea, 0.32 mM; Mitomycin C, 1 μg/mL). Scale bars, 25 μm. (D) Density distribution of 53BP1-mVenus foci numbers in single MERVL::tdTomato^−^ and MERVL::tdTomato^+^ cells (n = 108 for each subpopulation). (E) The time lag between asymmetric segregation of DNA damage and the point that MERVL::tdTomato intensity became different in sister lineages (n = 32). (F) Numbers of 53BP1-mVenus foci in 2C-entering and MERVL^−^ cells (n = 52 for each group) after DNA damage induction (Aphidicolin, 1.6 μM; Hydroxyurea, 0.08 mM; Mitomycin C, 0.25 μg/mL). (G) Asymmetric inheritance of 53BP1 foci between sister cells after DNA damage induction (Aphidicolin, 1.6 μM; Hydroxyurea, 0.08 mM; Mitomycin C, 0.25 μg/mL), plotted as the distribution of 53BP1-mVenus foci number of each sister divided by the sum of the pair (n = 55 pairs for each condition). *p* <0.0001. (H) Numbers of 53BP1 foci in MERVL^−^ (n = 110), 2C-death (n = 55) and 2C-survived daughter cells (n = 55) after DNA damage induction (Aphidicolin, 1.6 μM; Hydroxyurea, 0.08 mM; Mitomycin C, 0.25 μg/mL). (I) A schematic to depict the two correlation modes between asymmetric segregation of 53BP1 foci number and asymmetric division of MERVL^+^ cells after DNA damage induction (Aphidicolin, 1.6 μM; Hydroxyurea, 0.08 mM; Mitomycin C, 0.25 μg/mL). Left, Cells inheriting more 53BP1 foci numbers tend to adopt 2C-death fate; right, cells inheriting more 53BP1 foci numbers tend to adopt 2C-survived fate. The proportion of the two is shown in the pie chart. (n = 47 pairs for each situation, *p* = 0.011). (J) Numbers of 53BP1 foci in dying MERVL reporter-high cells (n = 30) and the few survived MERVL reporter-high cell (n = 35). Error bars indicate mean ± SEM (E, J). Mann–Whitney U test (F, H, J); permutation test (I); Kolmogorov-Smirnov test (G).

**Figure S4.**
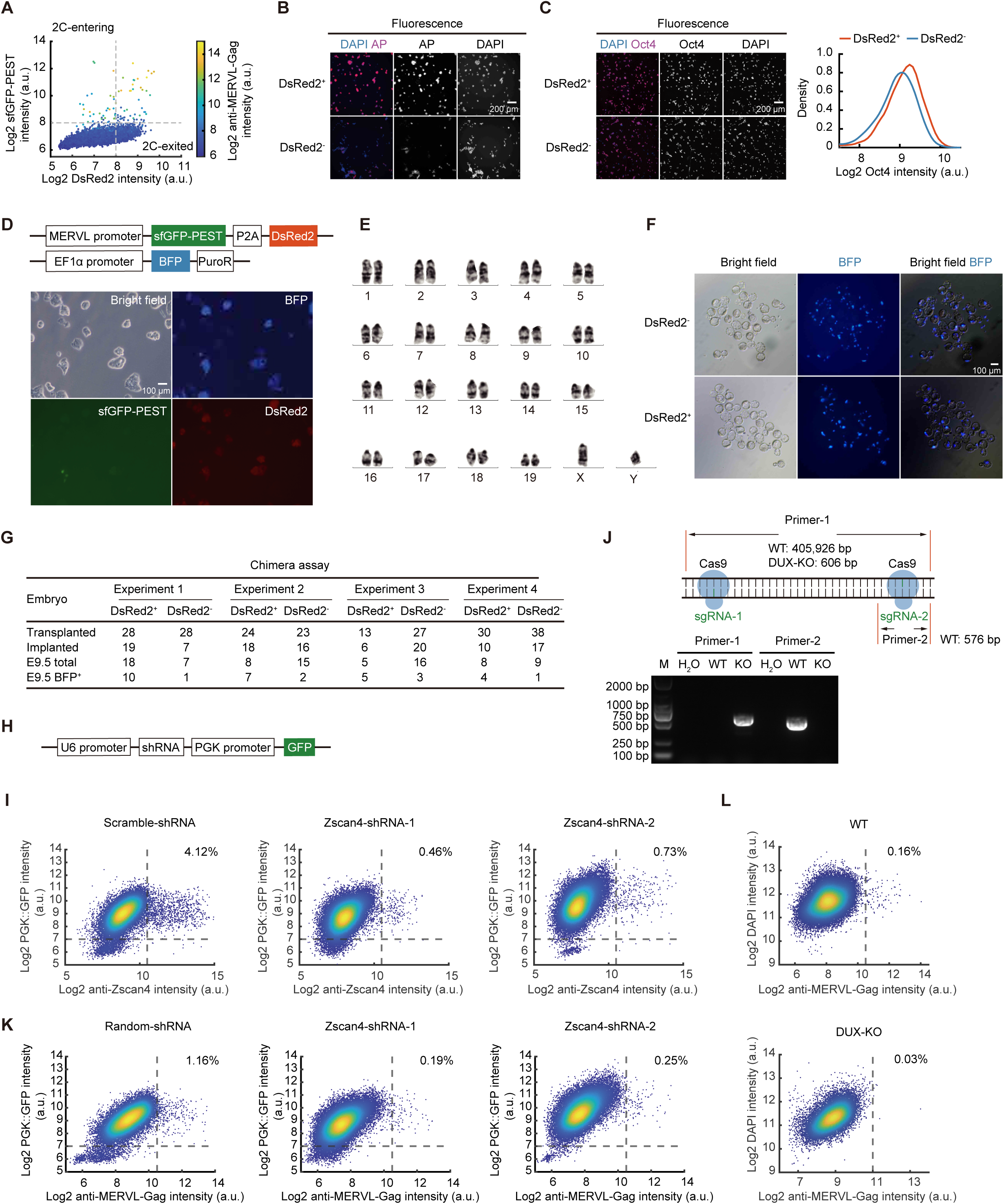
2C-exited cells show enhanced pluripotency, related to Figure 4. (A) Scatter plot of MERVL::sfGFP-PEST versus MERVL::DsRed2. The heat color indicates the intensity of MERVL-Gag antibody staining. MERVL-Gag antibody positive cells are mostly sfGFP^+^ cells, whereas DsRed2^+^ sfGFP^−^ cells are predominantly MERVL-Gag negative, suggesting these cells already exited the 2C-like state. (B) Representative fluorescent images of AP staining in sorted subpopulations in Figure 4C. Scale bar, 200 μm. (C) Oct4 antibody staining of sorted subpopulations in Figure 4C. Left, representative images; right, quantification (n = 5,656 for DsRed2^+^ sfGFP-PEST^−^ and n = 8,030 for DsRed2^−^ sfGFP-PEST^−^). Scale bar, 200 μm. (D) Top: design of the mESC line expressing MERVL::sfGFP-PEST-P2A-DsRed2 and EF1α-BFP constructs; bottom: representative images of the reporter cell line in GFP, DsRed2 and BFP channels. Scale bar, 100 μm. (E) Karyotype of the cell line in (D). (F) Microscope images of blastocysts cultured for one day after injection with sorted subpopulations in Figure 4C. Scale bar, 100 μm. (G) Numbers of transplanted, implanted, total and chimeric embryos at E9.5 in four independent experiments. (H) The construct used in the experiments of (J, L). (I) Quantification of Zscan4 immunofluorescence staining in the indicated mESCs. The percentages are quantified as the number of Zscan4^+^ PGK::GFP^+^ cells/PGK::GFP^+^ cells. PGK::GFP serves as a transfection control. (J) Genotype verification of the DUX knock-out (DUX-KO) C57 mESC line. (K) Quantification of MERVL-Gag immunofluorescence staining in the indicated mESCs. The percentages are quantified as the number of MERVL-Gag^+^ PGK::GFP^+^ cells /PGK::GFP^+^ cells. PGK::GFP serves as a transfection control. (L) Quantification of MERVL-Gag immunofluorescence staining in wild-type (WT, top) or DUX-KO (bottom) C57 mES cells.

## Video legends

**Video S1. MERVL::GFP^+^ cells undergo cell death, related to Figure 1**

**Video S2. Distinct cell fates of MERVL^+^ sister cells, related to Figure 1**

Cyan: mother cell; yellow: 2C-death cells; red: 2C-survived cells.

**Video S3. Asymmetric division of Zscan4^+^ cells, related to Figure 1**

Cyan: mother cell; yellow: 2C-death cells; red: 2C-survived cells.

**Video S4. Asymmetric segregation of 53BP1 foci in MERVL^+^ cells, related to Figure 2**

Red: mother cell; cyan: 2C-death cells; yellow: 2C-survived cells.

